# Pharmacologic and genetic inhibition of cholesterol esterification reduces tumour burden: a pan-cancer systematic review and meta-analysis of preclinical models

**DOI:** 10.1101/2021.06.12.448188

**Authors:** Alex Websdale, Yi Kiew, Philip Chalmers, Xinyu Chen, Giorgia Cioccoloni, Thomas A Hughes, Xinyu Luo, Rufaro Mwarzi, Marc Poirot, Hanne Røberg-Larsen, Ruoying Wu, Mengfan Xu, Michael A. Zulyniak, James L Thorne

**Affiliations:** School of Food Science and Nutrition, University of Leeds, Leeds, LS2 9JT, UK; School of Medicine, University of Leeds, Leeds, LS9 7TF, UK; Cancer Research Center of Toulouse, Inserm, CNRS, University of Toulouse, Toulouse, France; Department of Chemistry, University of Oslo, Norway

## Abstract

Cholesterol esterification proteins Sterol-O acyltransferases (SOAT) 1 and 2 are emerging prognostic markers in many cancers. These enzymes utilise fatty acids conjugated to coenzyme A to esterify cholesterol. Cholesterol esterification is tightly regulated and enables formation of lipid droplets that act as storage organelles for lipid soluble vitamins and minerals, and as cholesterol reservoirs. In cancer, this provides rapid access to cholesterol to maintain continual synthesis of the plasma membrane. In this systematic review and meta-analysis, we summarise the current depth of understanding of the role of this metabolic pathway in pan-cancer development. A systematic search of PubMed, Scopus, and Web of Science for preclinical studies identified eight studies where cholesteryl ester concentrations were compared between tumour and adjacent-normal tissue, and 24 studies where cholesterol esterification was blocked by pharmacological or genetic approaches. Tumour tissue had a significantly greater concentration of cholesteryl esters than non-tumour tissue (p<0.0001). Pharmacological or genetic inhibition of SOAT was associated with significantly smaller tumours of all types (p≤0.002). SOAT inhibition increased tumour apoptosis (p=0.007), CD8+ lymphocyte infiltration and cytotoxicity (p≤0.05), and reduced proliferation (p=0.0003) and metastasis (p<0.0001). Significant risk of publication bias was found and may have contributed to a 32% overestimation of the meta-analysed effect size was overestimated. Avasimibe, the most frequently used SOAT inhibitor, was effective at doses equivalent to those previously reported to be safe and tolerable in humans. This work indicates that SOAT inhibition should be explored in clinical trials as an adjunct to existing anti-neoplastic agents.

## 1. Introduction

Esterification is a tightly regulated component of cholesterol homeostasis and enables cholesterol packaging into lipid droplets. Intra-cellular storage of cholesterol allows ready access to meet the high demand for *de novo* plasma membrane synthesis during the rapid proliferation of cells, for example during tumour growth. Several diseases are linked to cholesterol esterification including a range of neurological conditions, lipid disorders, and cancer. The synthesis of cholesteryl esters (CE) is catalysed by Sterol O-acyltransferase 1 and 2 (SOAT1 and SOAT2), and Lecithin cholesterol acyltransferase (LCAT). SOAT1 and SOAT2, utilise fatty acid-coenzyme A conjugates to preferentially generate oleoyl (Fig1A), and linoleoyl or palmitoyl CEs (Fig1B), respectively, and coenzyme A as a biproduct [1]. LCAT utilises phosphatidylcholine (lecithin) to produce oleoyl CEs [2] but instead of producing coenzyme A as a biproduct, as is the case for SOAT1 and SOAT2, lysophosphatidylcholine is the biproduct (Fig1C). SOAT1 is ubiquitously expressed in tissues while SOAT2 is restricted to small intestines and the liver [3] and LCAT is expressed in the liver and secreted into circulation in lipoprotein complexes [4, 5].

**Figure 1.**
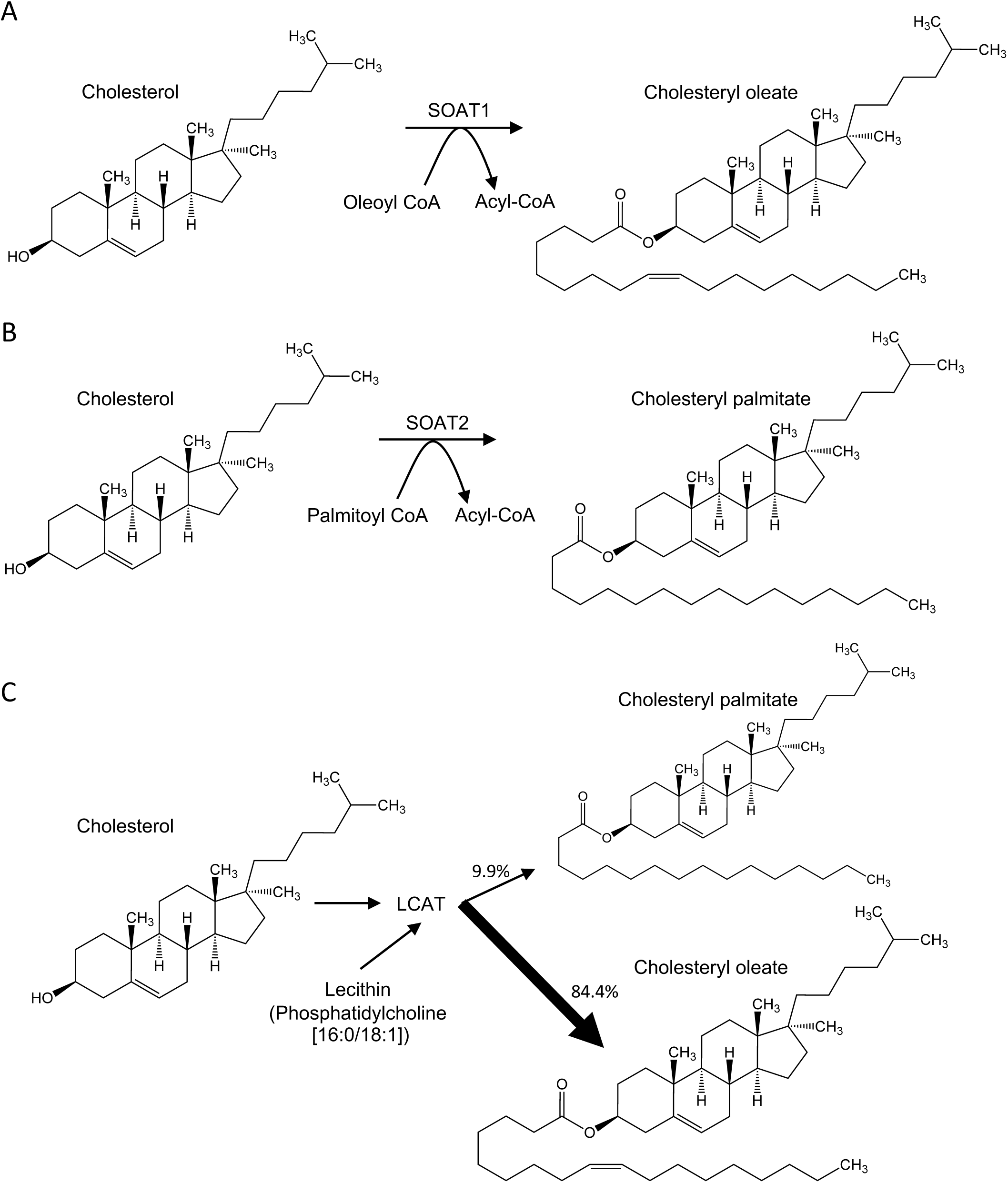
Mechanisms of cholesterol esterification. (A) The preferred substrates and products of SOAT1. (B) The preferred substrates and products of SOAT2. (C) The preferred substrates and products of LCAT. Reaction specificities of SOAT1 and SOAT2 were determined in SOAT1 or SOAT2 expressing H5 cells [1]. Reaction specificities for LCAT were determined using LCAT isolated from human serum [4].

Esterification of cholesterol in tumours is beneficial for cancer growth and studies of SOAT1, SOAT2 and LCAT in humans indicate this metabolic process is deregulated in cancer. Elevated SOAT1 and SOAT2 expression in tumours has been linked to higher grade of breast [6] and renal [7] cancer, respectively, and high expression SOAT1 has also been linked to poor prognosis for patients with liver [8], glioma [9], pancreatic [10] and adrenocortical [11] cancers. Furthermore, increased intracellular lipid droplet content, indicative of cholesterol esterification, is associated with reduced overall survival [12] and elevated cholesteryl oleoyl ester levels has been proposed as a prognostic biomarker for prostate cancer [13]. Conversely, high LCAT expression is associated with improved prognosis for liver cancer patients [14] and is often lower in liver cancer than normal tissue in humans [15] and rat models [16–18]. At the molecular level, cholesteryl esters promote cancer proliferation and invasiveness [19] and thus SOATs were considered as promising targets and the anticancer action of natural SOAT inhibitors such as auraptene and bryonolic acid was elucidated [20, 21]. Several small molecule inhibitors of cholesterol esterification have been explored in clinical trials for non-cancer related diseases providing an extensive understanding of their tolerability, toxicity and side-effect profiles. Avasimibe, first discovered in 1996 [22], is a dual SOAT1 and SOAT2 inhibitor [23, 24] and has been used in clinical trials for coronary atherosclerosis [25] and homozygous familial hypercholesterolemia [26]. Avasimin is human serum albumin encapsulated avasimibe that was developed to improve avasimibe solubility [27]. K-604 is a SOAT1 specific inhibitor [24] and has been tested for both safety and efficacy as a treatment against atherosclerosis (NCT00851500), however results from the trial have not been published. ATR-101 (Nevanimibe) is a SOAT1 specific inhibitor [28] that has been tested in a clinical trial against adrenocortical carcinoma (NCT01898715) [29] and Cushing’s syndrome (NCT03053271). Pactimibe also inhibits both SOAT1 and SOAT2 [23, 30], but a clinical trial (NCT00151788) administering 100 mg/day was terminated early due to a significant increase in major cardiovascular disease events [31]; pactimibe remains untested in pre-clinical cancer models. Drugs targeting CE synthesis that have been evaluated in clinical trials are summarised in Table 1.

**Table 1.** SOAT inhibitors assessed in clinical trials.

| Drug | Target | NCT | Reference | Condition or disease | Phase | Outcomes |
| --- | --- | --- | --- | --- | --- | --- |
| **Avasimibe (CI-1011)** | SOAT |  | [93] | Short-term safety |  | Avasimibe was tolerated at 500 mg daily for 8 weeks. Avasimibe induced reductions in triglycerides and VLDL cholesterol. |
| **Avasimibe (CI-1011)** | SOAT | NA | [25] | Atherosclerosis | NA | Avasimibe was tolerated at maximum dosage of 750 mg daily for 24 months. Avasimibe caused a moderate increase in LDL cholesterol and did not alter coronary atherosclerosis. |
| **Avasimibe (CI-1011)** | SOAT | NA | [26] | Homozygous familial hypercholesterolemia | NA | Avasimibe monotherapy was tolerated at 750 mg for 6-weeks. Avasimibe did not induce any significant lipid changes. |
| **K-604** | SOAT1 | NCT00851500 | Completed, no results published | Atherosclerosis | Phase 2 | NA |
| **Nevanimibe (ATR-101)** | SOAT1 | NCT01898715 | [94] | Adrenocortical carcinoma | Phase 1 | Nevanimibe was tolerated at up to 158.5 mg for 5 weeks. No tumour response to treatment at any dosages. |
| **Nevanimibe (ATR-101)** | SOAT1 | NCT02804178 | [95] | Congenital adrenal hyperplasia | Phase 2 | Nevanimibe was tolerated at 1000 mg twice daily for 2 weeks. Nevanimibe reduced 17-hydroxyprogesterone levels. |
| **Nevanimibe (ATR-101)** | SOAT1 | NCT03669549 | Terminated | Congenital adrenal hyperplasia | Phase 2 | NA |
| **Nevanimibe (ATR-101)** | SOAT1 | NCT03053271 | Terminated | Endogenous Cushing's syndrome | Phase 2 | NA |
| **Pactimibe (CS-505)** | SOAT | NCT00151788 | [96] | Familial hypercholesterolemia | Phase 2/3 | Pactimibe at a dosage of 100 mg increased low-density lipoprotein cholesterol. Pactimibe increased incidence of major cardiovascular events. |
| **Pactimibe (CS-505)** | SOAT | NCT00185042 | Completed, no results | Coronary artery disease | Phase 2 | NA |
| **Pactimibe (CS-505)** | SOAT | NCT00185146 | Completed, no results published | Atherosclerosis | Phase 2 | NA |

There is a significant body of research investigating cholesterol esterification in pre-clinical cancer models. Many of these studies utilise pharmacological or genetic inhibitors of SOAT to provide insight into the cellular and molecular role of the enzyme and propose repurposing SOAT inhibitors as cancer therapies. However, the pharmacological compounds remain underexplored in the clinical cancer setting and may be suitable for repurposing. This systematic review and meta-analysis summarises the evidence regarding cholesterol esterification enzymes as therapeutic targets in cancer and details the range of pharmacological approaches that are closest to clinical translation.

## 2. Methods

### 2.1 Search strategy

The search strategy was applied to PubMed, SCOPUS, Web of Science and the Cochrane Library, and records were retrieved up until April 2021; the strategy is registered in the PROSPERO database (CRD42020202409) with the following modifications: records evaluating sulfation and associated enzymes were excluded from the study.

### 2.2 Study selection

Titles and abstracts were screened against inclusion criteria; (i) original research, (ii) investigated cancer, (iii) assessed an *in vivo* pre-clinical animal model and (iv) modulated cholesterol esterification. Each abstract was assessed by two independent assessors and discrepancies resolved by a third member of the research team. All publications that satisfied the above criteria were included for qualitative assessment.

### 2.3 Data extraction

Publication that reported adequate data for quantitative assessment were included in the meta-analyses. Data were extracted in duplicate by two independent assessors and discrepancies resolved as a team. Mean values and measures of variance were extracted. Only data from test groups assessing either CE concentration, or enzyme expression/activity were extracted and studies reporting combination therapies or other enzymes such as SULT2b were excluded at this point. Where data was not available in the text, data was extracted from appropriate figures using WebPlotDigitizer (v4.2) by two independent assessors. Data regarding animals, study design, mechanism of SOAT1 disruption, cancer type and outcomes assessed were extracted.

### 2.4 Statistical analysis

Review Manager version 5.4 (The Nordic Cochrane Centre, 2014) was used to perform meta-analyses. Where more than one treatment dose was measured in comparison to the control, the largest dose was used for meta-analysis. Where studies reported data as fold change relative to the starting volume, fold changes were normalised to tumour size at the initiation of the experiment by us to standardise study data. Tumour sizes were standardised to cm^3^ across studies. Mean difference was used where appropriate but where differed from the same outcome standardised mean difference (SMD) was used. SMD effect size is interpreted as mean difference relative to the variance observed in the comparison. Random effects model was used due to the anticipation of heterogeneity between studies due to expected differences in cancers assessed, animal models and mechanisms used to disrupt SOAT1 [32]. Heterogeneity was assessed using I^2^, with an I^2^ value >75% used as a marker for high heterogeneity between studies due to the anticipated large variation between study design for animal studies [33]. Evidence of publication bias was examined using funnel plots.

### 2.5 Publication Bias

When publication bias was apparent within funnel plots, a corrective overestimation value was determined with Duval and Tweedle’s trim and fill method using Comprehensive Meta Analyst version 3 (Biostat inc., 2014). In cases where analyses exhibited an I^2^ value <25% or >75%, reasoning behind their heterogeneity, or lack of, was discussed.

### 2.6 Risk of bias

Risk of bias (ROB) was adapted from Cioccoloni et al., and allowed assessment of bias in experimental design, animal experiments, and immunoblotting [33]. Guidelines published in British Journal of Pharmacology [34, 35] and SYRCLE [36] were closely followed.

## 3. Results

### 3.1 Systematic search

#### 3.1.1 Records returned

A systematic search strategy was applied to multiple databases, PubMed, SCOPUS, Web of Science, and the Cochrane Library, returning 847, 970, 847 and 20 records respectively; four additional records were identified during background reading. Following removal of duplicates there were 1543 unique records for screening. Abstract screening returned 76 records that had evaluated inhibition of SOAT1, SOAT2, or LCAT (n=43) or where intra/inter-tumoural CE concentrations were assessed (n=41). After full text screening, 24 records assessing pharmacological or genetic inhibition of cholesterol esterification enzymes were suitable for both qualitative and quantitative analyses. Thirteen studies on CE tumour concentrations were suitable for qualitative synthesis, of which ten were suitable for quantitative analysis. This information is summarised in Fig2A.

**Figure 2.**
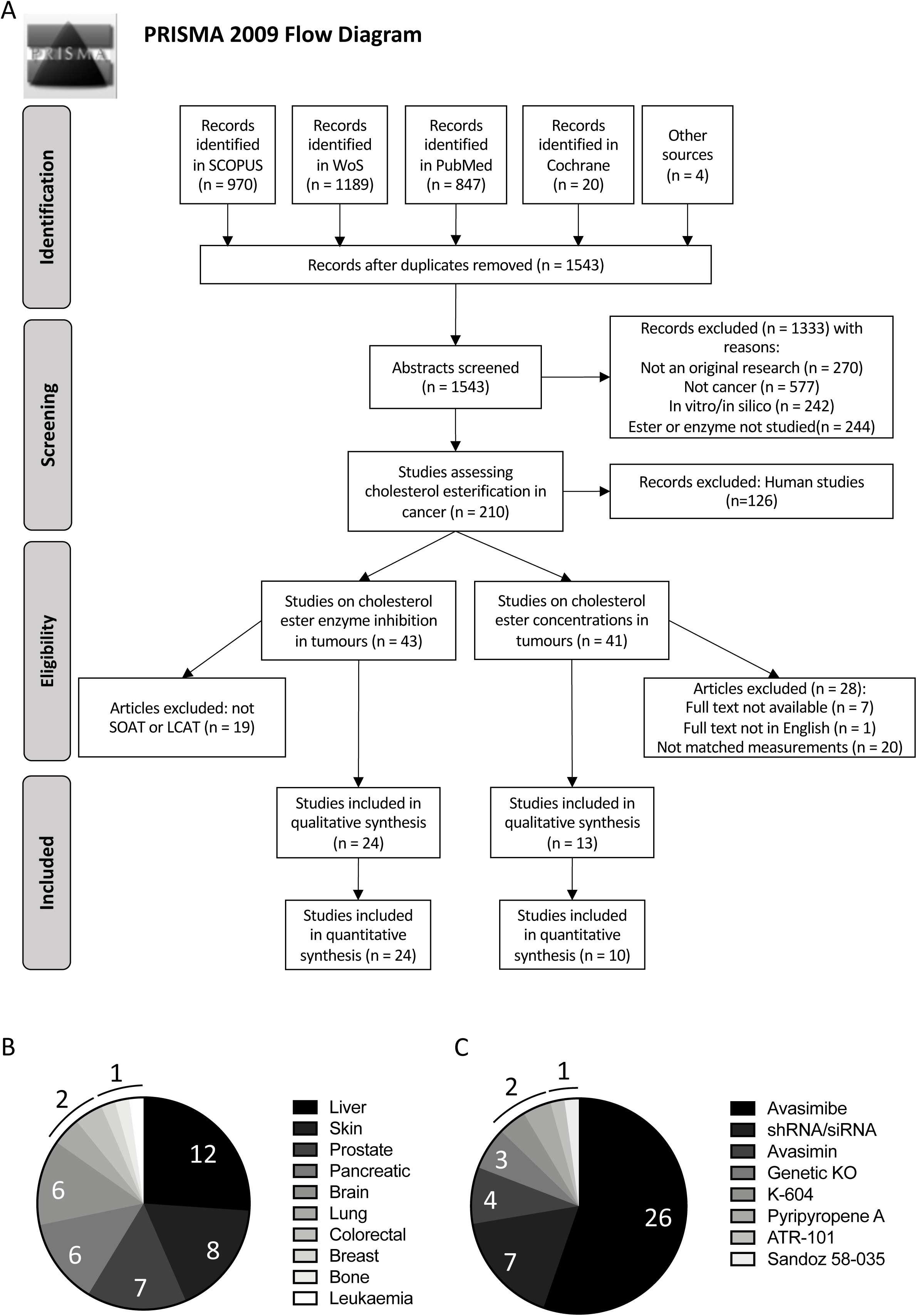
Study discovery and distribution. (A) PRISMA flow diagram showing searching, screening, eligibility and inclusion process. (B) Spread of papers assessing different cancer types in SOAT inhibition studies. (C) Spread of papers assessing different SOAT inhibiting treatments in SOAT inhibition studies.

#### 3.1.2 Cancer sites

All 46 comparisons within the 24 studies that were included in quantitative analysis of SOAT inhibition were mouse xenograft or allograft models assessing SOAT1 and/or SOAT2 inhibition. Of the 46 comparisons, 12 evaluated liver cancer, eight on skin cancer, seven on prostate cancer, six on pancreatic cancer, six on brain cancer, two on lung cancer, two on colorectal cancer, while breast cancer, bone cancer and leukaemia were each studied once (Fig2B). All comparisons that assessed SOAT1/2 inhibition were xenograft models. We found no records pertaining to LCAT inhibition or activation meeting our search criteria. Of the ten studies included in the quantitative analysis of cholesterol ester concentration in tumour and matched or non-matched normal tissue, six comparisons assessed liver cancer, two assessed testicular cancer, and one of each assessed breast, pancreatic and renal cancers. Seven were xenograft models, three were mutagen induced models and one was a radiation induced model of cancer.

#### 3.1.3 Interventions and dosing

Five small molecule inhibitors, RNAi, and genetic knock-out were used across the studies. Avasimibe was the most commonly used drug and was administered at between 2 to 30 mg/kg, typically at 15 mg/kg (16 times across 11 studies) but lower (2 mg/kg one study; 7.5 mg/kg three studies) and higher (30 mg/kg two studies) concentrations were evaluated. Avasimin was used in four comparisons across two studies (75 mg/kg) with or without supplementation with 7.5 mg/kg avasimibe. K-604 was used twice in one study (30 μg/cm^3^ tumour); ATR-101 also once (1mg/g chow); Sandoz 58-035 (a dual SOAT1/2 inhibitor [37]) also once (15 mg/kg). Pyripyropene A, a SOAT2 specific inhibitor, was used in one study, twice. Pre-treatment of cancer cells with shRNA or siRNA before grafting was the second most common intervention targeting SOAT1, after avasimibe treatment. Three comparisons assessed genetic knockout of SOAT1, with one performing SOAT1 knockout in the animals’ T-cells (Fig2C). Pre-treatment with siRNA against SOAT1 was performed in either cancer cells across six comparisons or CAR T-cells in one comparison. Pre-treatment of cancer cells with siRNA against SOAT2 was assessed in one comparison. Importantly, SOAT is also known as acyl-coenzyme A:cholesterol acyltransferase (ACAT) and has been confused in the literature previously with acetyl-coenzyme A acetyltransferase, also referred to as ACAT. To add further confusion, acetyl-coenzyme A acetyltransferase also has two isoforms, 1 and 2, mimicking that of SOAT1 and SOAT2. This confusion has led the use of improper reagents [38] and studies otherwise meeting our search criteria were excluded from our analyses for this reason. Several studies did not consider avasimibe’s broadly equivalent IC_50_ against both SOAT1 and SOAT2 (Table 2) and reports that their data are SOAT1 specific is erroneous [23, 24]. Only one study tested SOAT2 specific inhibition using either siRNA or pyripyropene A.

**Table 2.**
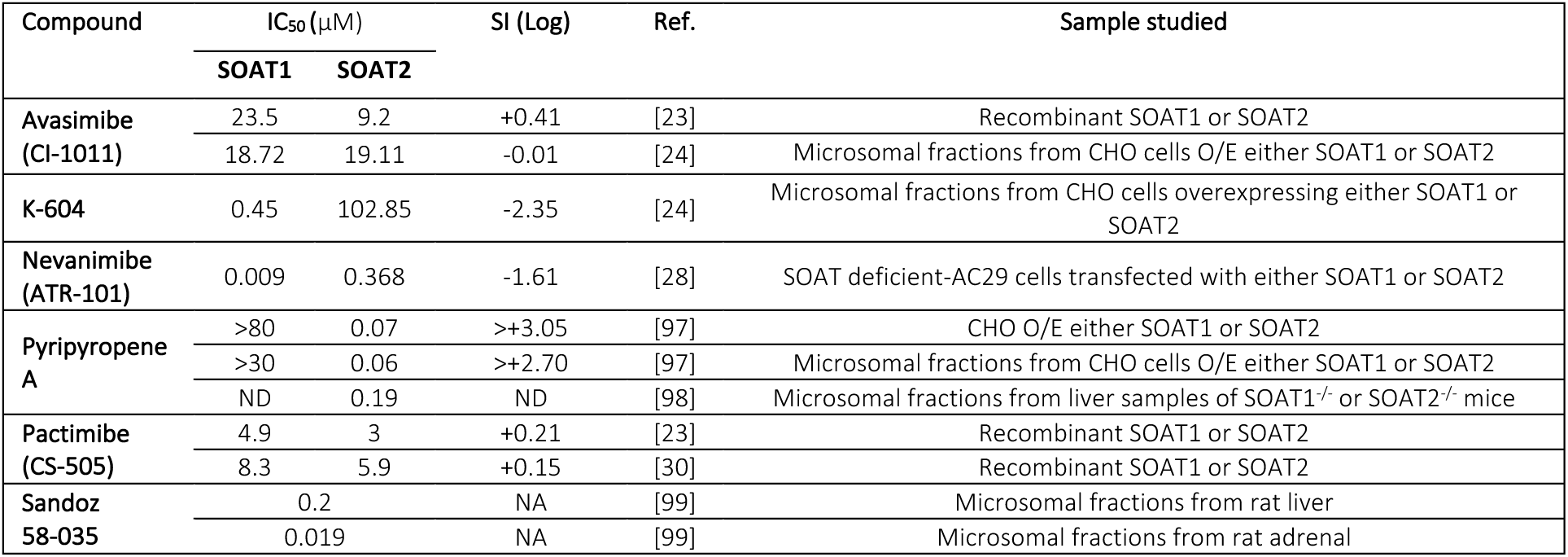
IC50 of SOAT inhibitors assessed in clinical or pre-clinical studies. Preferred isoform is indicated in bold if greater than 10-fold difference in IC_50_ has been reported. *SI (log) is log(IC_50_ for SOAT1/IC_50_ for SOAT2).

Drugs were administered by intraperitoneal injection (IP) in ten studies, intravenous (IV) in three studies, per oral (PO) in three studies, intragastric administration (IG) in two studies, and intratumoural (IT) and subcutaneously in one study each. Although avasimibe is an orally bioavailable drug [22], only two studies [39, 40] assessed tumour size following PO administration. Through this route 30 mg/kg avasimibe was effective against the bone cancer model, U2OS xenograft, leading to a 90% reduction in tumour volume, but a 15 mg/kg dose against the prostate cancer PC3 xenograft model was not effective.

### 3.2 Cholesterol esters are concentrated in tumour tissue

Eight studies compared CE concentrations in tumour and normal tissue from the same animals. These were largely evaluating liver cancer (n=6), with single studies each of testicular cell and renal cell carcinoma. CE concentrations were significantly higher in tumour tissue compared to non-tumour tissue from the same animal (SMD = 1.29; 95% CI: 0.68 to 1.90; I^2^ = 31%; p < 0.0001; Fig3A). Van Heushen et al. found CE concentrations were no different between microsomal fractions derived from xenograft and non-tumour tissue [41]. Harry et al., found CE increased in each of three different hepatocellular carcinoma xenograft models (Table 3) [42] but this was not included in our meta-analysis as SD were not reported. Surprisingly, when comparing tumour tissue from tumour-bearing animals with normal tissue from control animals there was no significant difference in CE concentration (Fig3B; p > 0.05).

**Figure 3.**
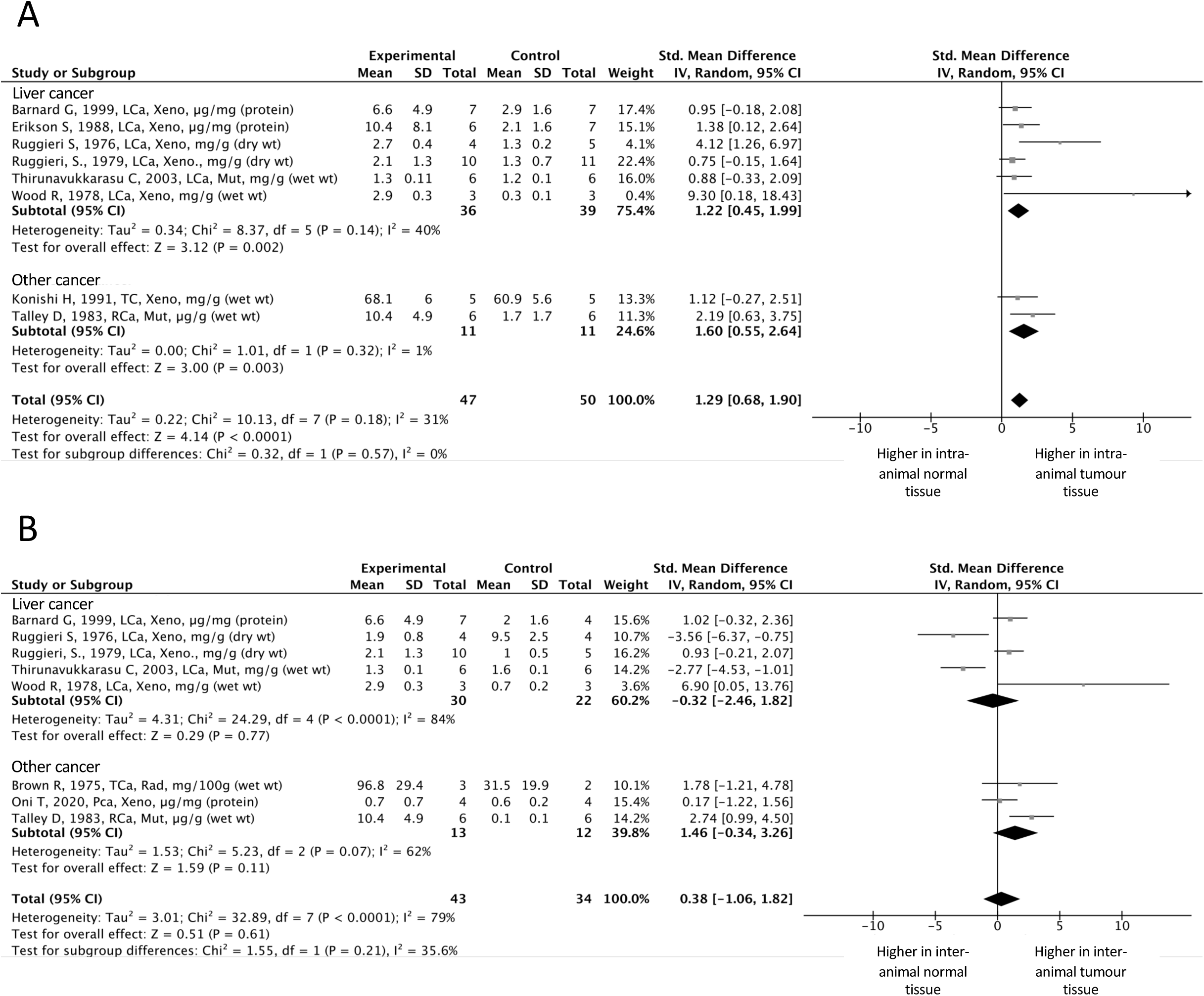
Cholesteryl ester concentration in tumour tissue and matched-normal tissue from control littermates. (A) Cholesteryl ester concentration in tumour tissue and matched normal tissue from the same mouse. (B) Cholesteryl ester concentration in tumour tissue and matched-normal tissue from control littermates. Differences in cholesterol ester concentration between tissues is represented as a standardised mean difference.

**Table 3.**
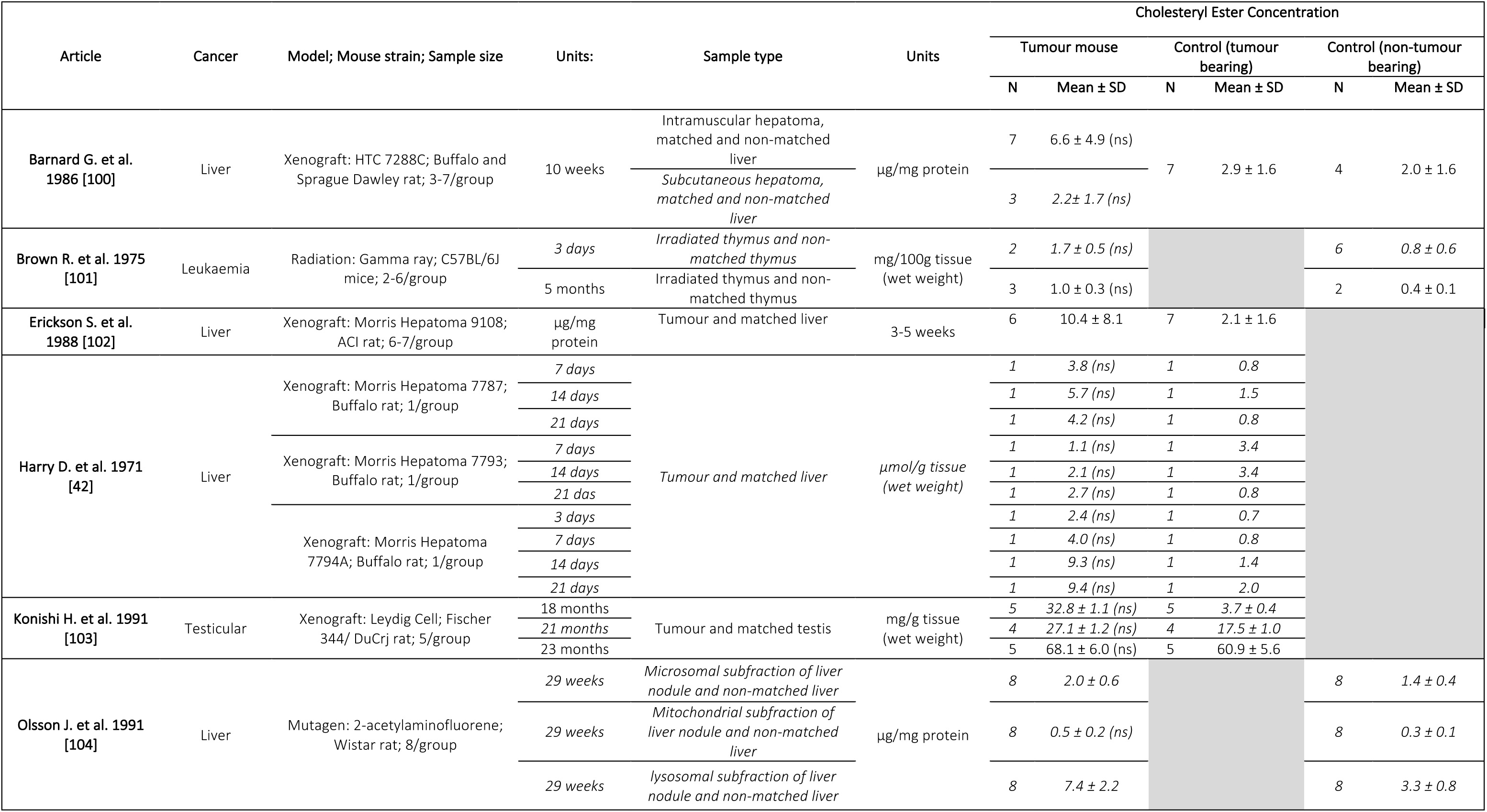
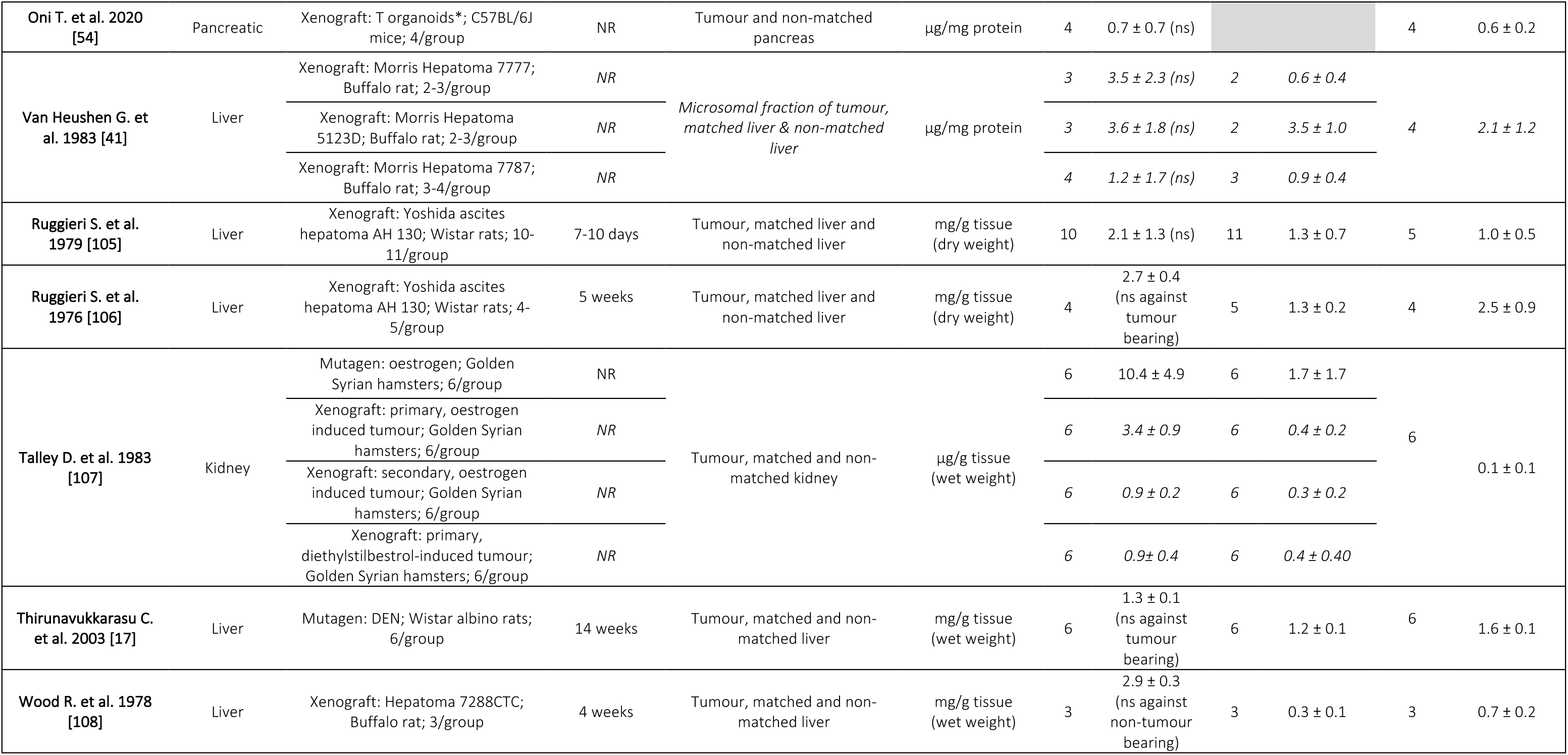
Summary of extracted data from cholesteryl ester measurement studies. Italic entries were not included in meta-analysis to avoid double counting of controls. Abbreviations: DEN = Diethylnitrosamine, T organoids = Xenografted tumour cells from KrasLSL-G12D/+; Trp53LSL-R172H/+; Pdx1-Cre mouse tumour. NR = not recorded.

### 3.3 SOAT promotes tumour growth

We next evaluated the impact of inhibiting cholesterol esterification enzyme expression or activity on tumour development. Twenty-four studies reported 40 comparisons of SOAT inhibition versus control treatment. Twenty-seven out of 40 comparisons found that tumours were significantly smaller after SOAT inhibition or knock-down compared to controls. Our meta-analysis (40 comparisons, total number of animals = 555) demonstrated that impairment of SOAT1 activity and/or expression is strongly associated with reduced tumour size (Fig4). We found sufficient studies to analyse separately size of brain, liver, pancreas, prostate, and skin cancer. Several other studies assessing other cancers were identified but not in sufficient numbers for individual analyses. These were instead grouped as ‘other cancers’.

**Figure 4.**
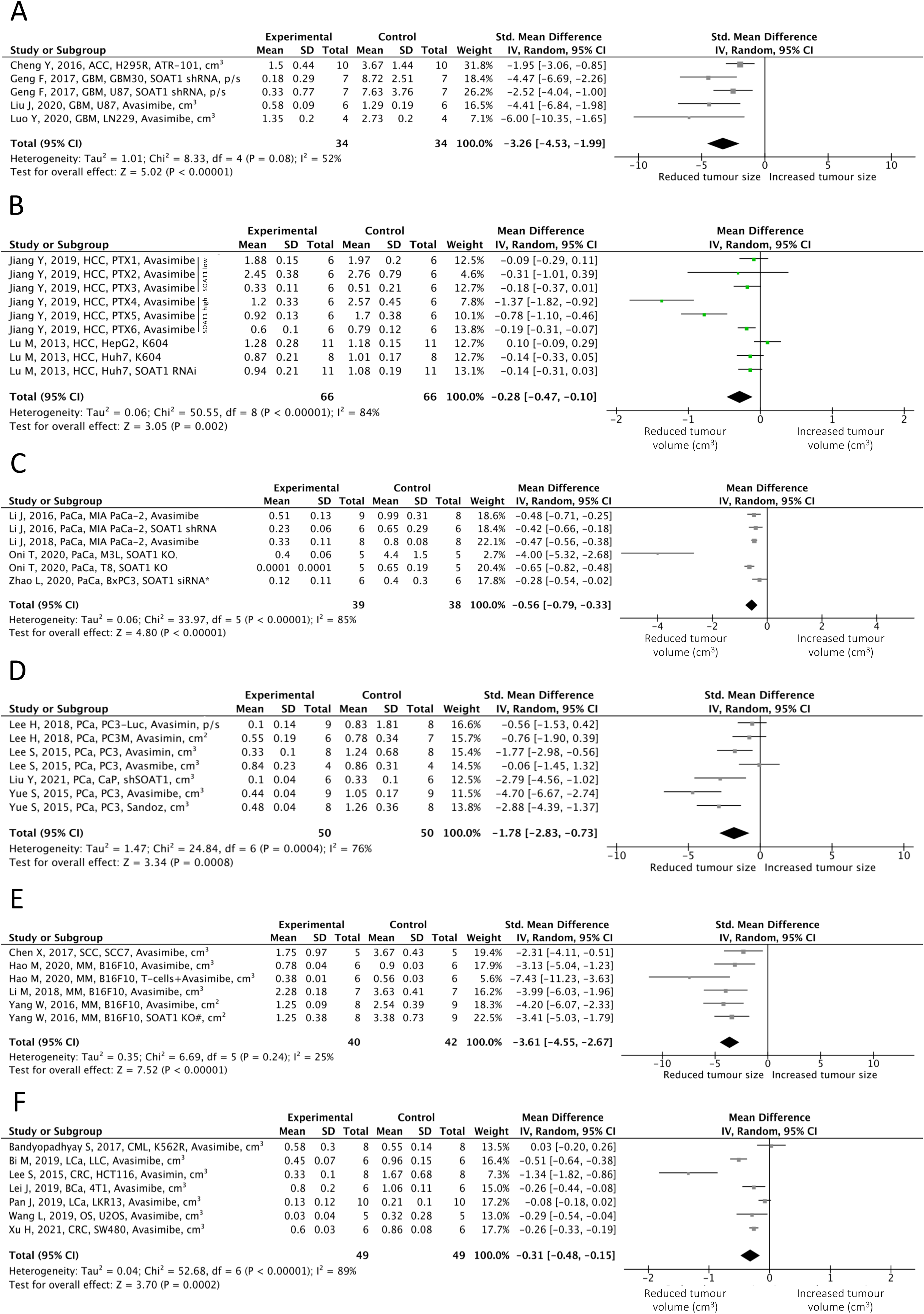
Change in tumour size following disruption of SOAT1. (A) Standardised mean difference in brain cancers. (B) Mean difference (mm^3^) in liver cancers. (C) Mean difference (mm^3^) in pancreatic cancers. (D) Standardised mean difference in prostate cancers. (E) Standardised mean difference in skin cancer. (F) Mean difference (mm^3^) in other cancers. * denotes modifications localized to CAR T-cells. # denotes modifications localized to T-cells.

#### 3.3.1 Brain cancer

Our systematic review identified four studies that explored SOAT inhibition in two brain cancer subtypes, glioblastoma [12, 43, 44] and adrenocortical cancer [45]. Glioblastoma is the most common primary malignant brain tumour, accounting for 48% of cases [46]. Adrenocortical carcinoma is a rare malignancy, with equally poor disease-free survival rates [47]. In all studies, SOAT1 inhibition led to a reduction in tumour size measured as either volume (cm^3^) or radiance (units of photons/seconds/cm^2^/units of solid angle or steradian, abbreviated to p/s) (SMD = −3.26; 95% CI: −4.53 to −1.99; I^2^ = 52%; p < 0.00001; Fig4A). In the U87 glioblastoma model, growth of xenografted cells was reduced using siSOAT1 [12] and avasimibe [43]. Liu et al. tested two doses of avasimibe (15 mg/kg and 30 mg/kg), but no dose response was observed with respect to either tumour volume or weight (Table 4). Avasimibe also impaired growth of LN229 xenografts, another glioblastoma model. Here the authors provided evidence that loss of the long non-coding RNA linc00339 mediated avasimibe’s anti-tumour effects; linc00339 overexpression prevented the avasimibe-mediated growth inhibition (Table 4) [44]. In adrenocortical brain cancer, PO administration of ATR-101 was associated with significantly smaller H295R xenografts compared to controls (Table 4) [45].

**Table 4.**
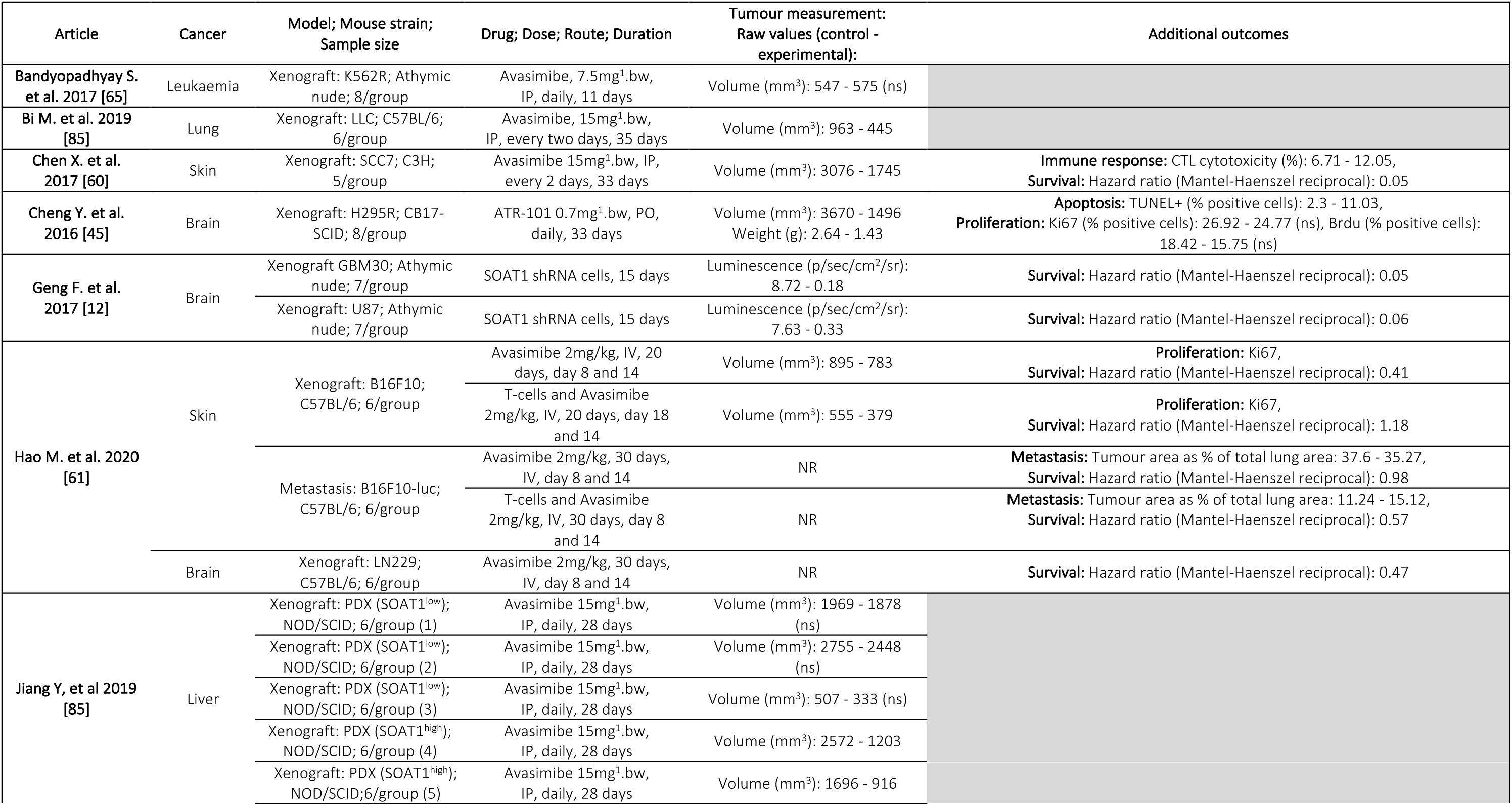
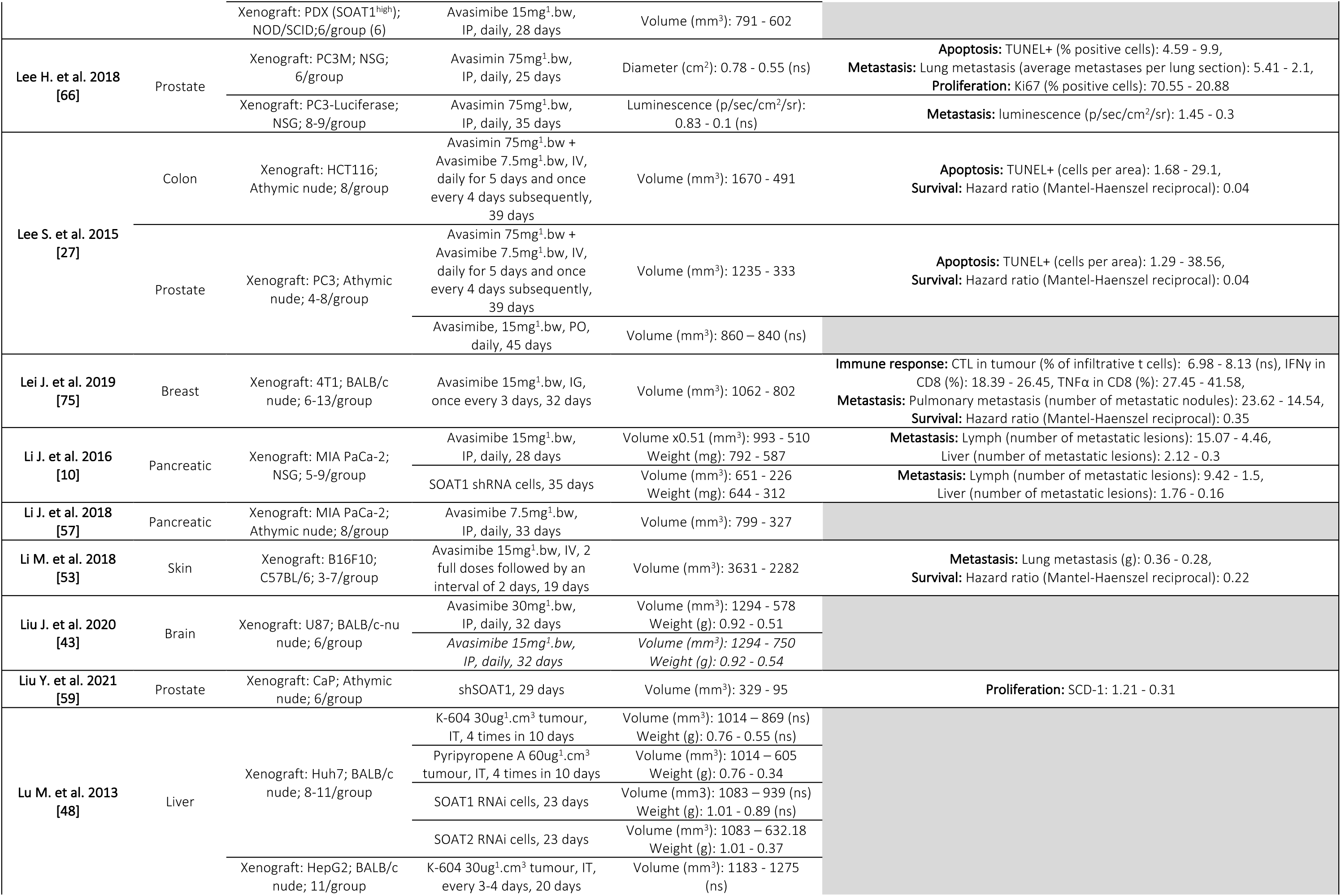
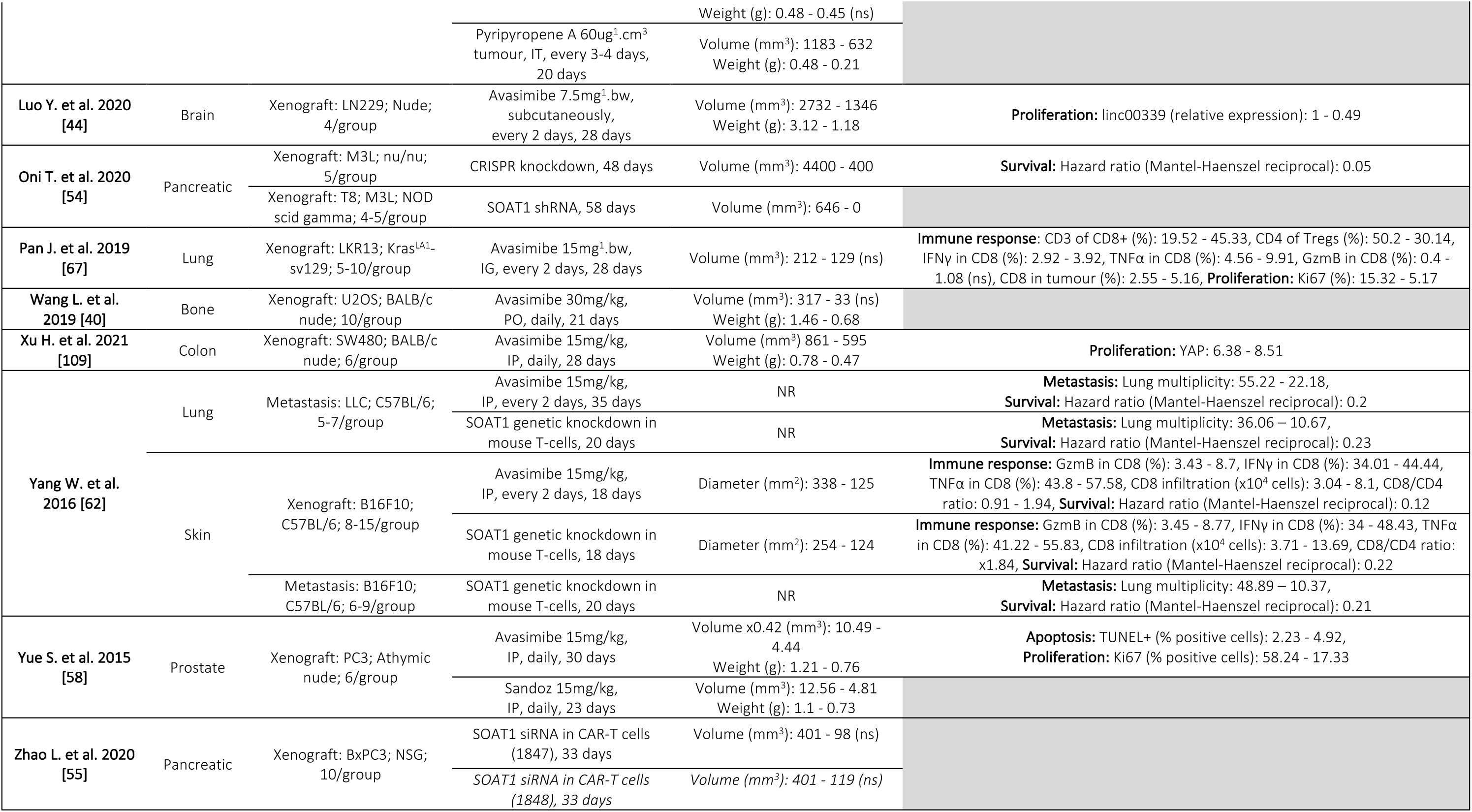
Summary of extracted data from SOAT1/2 inhibition studies. Italic entries were not included in meta-analysis. Abbreviations: bw = body weight, CAR-T = chimeric antigen receptor T, con. = control, CTL = cytotoxic t lymphocyte, exp. = experimental, GzmB = granzyme b, IFNγ = interferon gamma, IG = intragastric administration, IP = intraperitoneally, IV = intravenously, IT = intratumourally, NR = not recorded, ns = not significant, PDX = patient derived xenograft, SR = units of solid angle or steradian, TNFα = tumour necrosis factor.

#### 3.3.2 Liver cancer

We identified nine experiments from two publications suitable for inclusion in quantitative analysis of SOAT inhibition in liver cancers [8, 48]. Liver cancer diagnoses, of which 80-90% are hepatocellular carcinoma, are the third most prevalent cause of cancer death in the world [49]. SOAT inhibition was associated with significantly smaller liver tumours (MD = −0.28; 95% CI: −0.47 to −0.1; I^2^ = 84%; p = 0.002; Fig4B). SOAT1 expression was measured in patient derived xenografts (PDX), and interestingly, avasimibe was most effective at reducing tumour volume in those expressing high levels of SOAT1; in PDXs with low SOAT1 expression tumour response was modest or absent [8]. In other liver models, notably xenograft of Huh7 or HepG2, inhibition of SOAT2 but not SOAT1 was associated with smaller tumour volumes. Intratumoural injection of K-604, a SOAT1 selective inhibitor, was not associated with smaller tumours, nor was siSOAT1 pre-treatment of Huh7 prior to implantation [48]. Instead siSOAT2 of the Huh7 cells before transplant led to significantly smaller tumours than controls (Table 4). Furthermore, Huh7 and HepG2 xenografts treated with Pyripyropene A, a selective SOAT2 inhibitor, were also smaller than control xenografts [48]. This may be explained by expression levels; SOAT2 is expressed at higher levels than SOAT1 in HepG2 cells according to The Protein Atlas (Huh7 not available) [50, 51] and SOAT2 has been reported as frequently upregulated in hepatocellular carcinoma [52].

#### 3.3.3 Pancreatic cancer

Our systematic review identified six experiments performed in four publications [10, 53–55], all of which assessed tumour volume in pancreatic cancer. Pancreatic cancer is the 14^th^ most common cancer in the world [49], with a 5-year survival rate of just 7% [56]. The mean difference between treatment and control groups was calculated and tumours in the SOAT inhibition groups were on average more than 0.5 cm^3^ smaller than control tumours (MD = - 0.56; 95% CI: −0.79 to −0.33; I^2^ = 85%; p < 0.0001; Fig4C). Zhao et al. produced chimeric antigen receptor T-cell (CAR-T) variants and injected into BxPC3 xenografts. In two siSOAT1 knockdown experiments, tumour growth was slower relative to control, yet there was no change in CAR-T infiltration into the tumour. The authors concluded SOAT1 is required for CAR-T anti-tumour cytotoxicity but not tumour homing. Li et al. examined direct shRNA knockdown of SOAT1 in MIA PaCa-2 cells and reported 0.5cm^3^ smaller tumours relative to controls. In one instance, pre-treatment of xenografted cells with shSOAT1 fully suppressed tumour formation [54]. As no mean or SD was reported within this comparison due to absence of tumour at the final timepoint, a measure near zero was imputed (1×10^-4^ cm^3^) for the mean and SD to allow use in the meta-analysis. Avasimibe has also been tested in the pancreatic setting, and interestingly was found to be more effective at impairing MIA PaCa-2 xenograft growth in a tumour placed subcutaneously [57] as opposed to within the pancreas [10].

#### 3.3.4 Prostate cancer

Prostate cancer is the most common cancer affecting men in the world; nearly 1.5 million new cases are diagnosed each year [49]. Growth of preclinical prostate cancer models are significantly impaired by inhibition of cholesterol esterification (SMD = −1.78; 95% CI: −2.83 to −0.73; I^2^ = 76%; p = 0.0008; Fig4D). Three drugs were examined in three studies. The efficacy of PO avasimibe was lower than IV avasimin [39], and Sandoz was less efficacious than avasimibe [58]. Prostate cancer cells have recently been shown to be highly sensitive to loss of SOAT1. CaP cells that were pre-treated with shSOAT1 prior to xenografting grew into significantly smaller tumours than their control counterparts [59].

#### 3.3.5 Skin cancer

Skin cancers (melanoma and non-melanoma) are the third most prevalent cancer type in the world [49]. Tumour burden was significantly lower in models of skin cancer that had been treated with SOAT1 inhibitors across all four studies relevant for quantitative analysis [53, 60–62] (SMD = −3.61; 95% CI: −4.55 to −2.67; I^2^ = 25%; p < 0.00001; Fig4E). SOAT1 was genetically knocked out of the T-cells of mice rather than in the implanted B16F10 cells and interestingly produced a similar standardised mean difference in tumour volume to systemic avasimibe treatment [62]. Furthermore, the introduction of T-cells and avasimibe to lymphodepleted mice led to significantly smaller tumours than treatment with avasimibe alone [61]. Interestingly, animals from this study were treated with just 2 mg/kg avasimibe, considerably lower than the doses we found reported in other studies of skin, or any other cancer type.

#### 3.3.6 Other cancers

Seven other studies measured five other cancer types, finding tumours to be significantly smaller after systemic SOAT inhibition (p = 0.0002; Fig4F). Chronic myelogenous leukaemia (CML) was the only tumour type that did not respond to SOAT inhibition. Resistance to SOAT inhibition may be driven through the BCR-ABL translocation, which is very common in CML [63]. The BCR-ABL fusion activates multiple oncogenic signalling pathways including MAPK, AKT and MYC [64]. Interestingly, avasimibe treatment does decrease MAPK signalling, but changes in other pathways have not been reported [65].

### 3.4 SOAT expression is associated with enhancement of cancer hallmarks

#### 3.4.1 Sustained proliferative signalling

Tumour proliferative index provides information regarding the rate of tumour growth and can be measured by expression of Ki67 or PCNA, which are components of the cell cycle machinery, or via incorporation of synthetic nucleosides such as BrdU, which marks de novo DNA synthesis. SOAT1 inhibition was associated with significantly lower Ki67 positivity in cancer cells in four out of five studies (MD = −14.43; 95% CI: −22.32 to −6.55; I^2^ = 98%; p = 0.0003; Fig5A) that included xenograft models of PC3 [27, 66] and LKR13 [67] cells and allograft models of B16F10 [61] cells treated with avasimibe or avasimin. Surprisingly, in the B16F10 allograft model treated with both avasimibe and T-cells, there is an increase in Ki67+ cancer cells. However, this treatment still induced a significant reduction in tumour volume, suggesting other mechanisms may have mediated tumour destruction [61]. ATR-101 did not alter Ki67 expression or BrdU incorporation in H295R xenografts despite this being associated with reduced tumour size [45] (Table 4).

**Figure 5.**
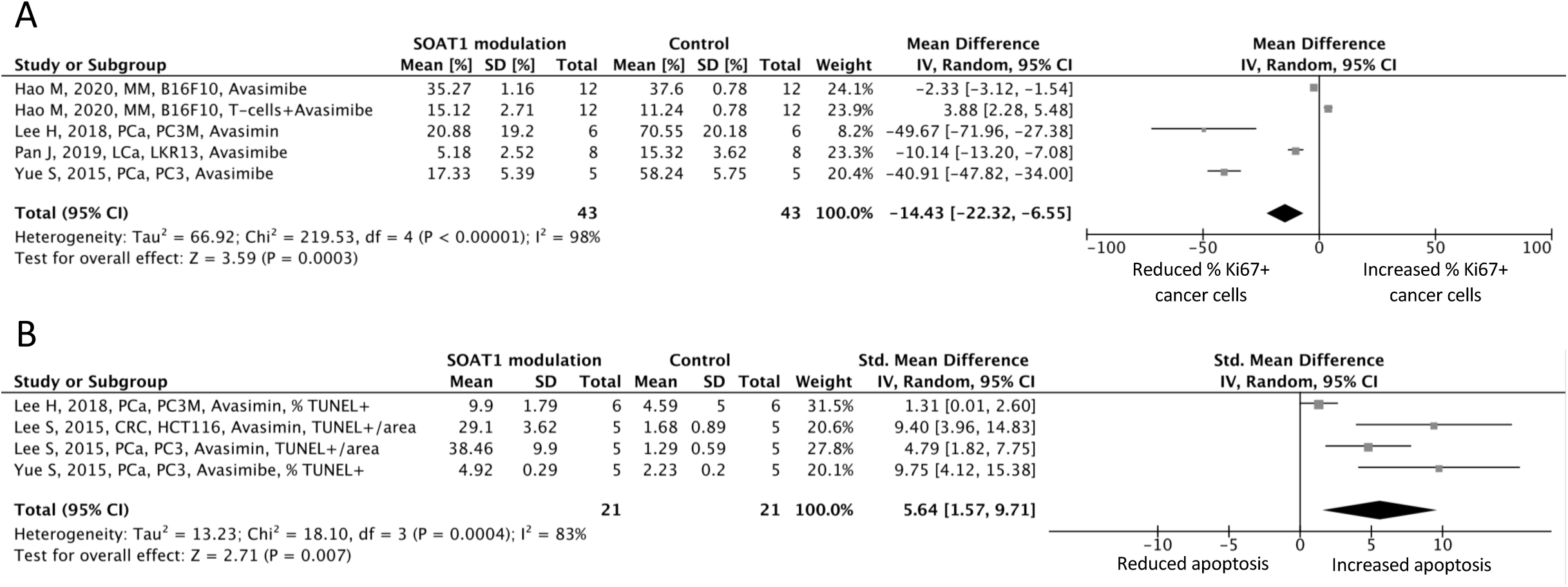
Forest plots showing changes in apoptosis and proliferation. (A) Mean difference (percentage Ki67+ cells) between experimental and control groups in tumour expression of Ki67. (B) Standardised mean difference between experimental and control groups in apoptotic cells in the tumour as measured by TUNEL+ stain assay.

#### 3.4.2 Resisting cell death

The ability of cancers to resist apoptosis enhances tumour growth. Commonly, cell death is assessed through apoptosis assays such as the TUNEL+ assay, expression of apoptosis mediating proteins, or mitochondrial function assays. Across four comparisons from three studies, TUNEL+ staining was significantly enhanced by avasimibe or avasimin (SMD = 5.64; 95% CI: 1.57 to 9.71; I^2^ = 83%; p = 0.007; Fig5B) in PC3 [27, 39, 66], PC3M [39], HCT116 [39] xenograft models. The same was found in H295R xenografts treated with ATR-101, suggesting that increased apoptosis rather than reduced proliferation is driving reduced tumour volume in this model [45]. Free cholesterol levels were also elevated in PC3 and HCT116 xenograft models undergoing apoptosis after avasimibe exposure [39].

#### 3.4.3 Evasion of immune detection

The immune system’s anti-tumour response can be activated following detection of tumour antigens by CD8+ T-cells. High levels of cytotoxic T-cell infiltration into tumours indicates a good prognosis for patients with breast [68], colorectal [69], lung [70], skin [71] and prostate cancer [72]. However, if the invaded T-cell population is anergic they are unable to mount a sufficient cytotoxic response and anti-tumour efficacy is severely reduced [73]. Our meta-analysis indicated that inhibition of SOAT1 was associated with increased CD3+CD8+ and CD8+ cytotoxic T lymphocytes (CTL) infiltration into the tumour (SMD = 1.12; 95% CI: 0.46 to 1.77; I^2^ = 0%; p = 0.0009; Fig6A). Not only did avasimibe treatment stimulate a time-dependent increase in CTL infiltration but the drug was also shown to impair efficiency of the immunosuppressive tumour environment through a decrease in the tumour’s CD4+ Tregs count [67]. As Tregs suppress CD8+ cell proliferation [74], this may explain why CD8+ cell infiltration increased. Treg infiltration was unaffected during a T-cell specific knockout of SOAT1 in a melanoma xenograft model [62] but CD8+ infiltration into tumour was induced at similar levels to avasimibe treatment, suggesting that disruption of cholesterol esterification in CD8+ cells alone is enough to induce increased infiltration, independently of systemic SOAT1 inhibition and the CD4+ Treg population.

**Figure 6.**
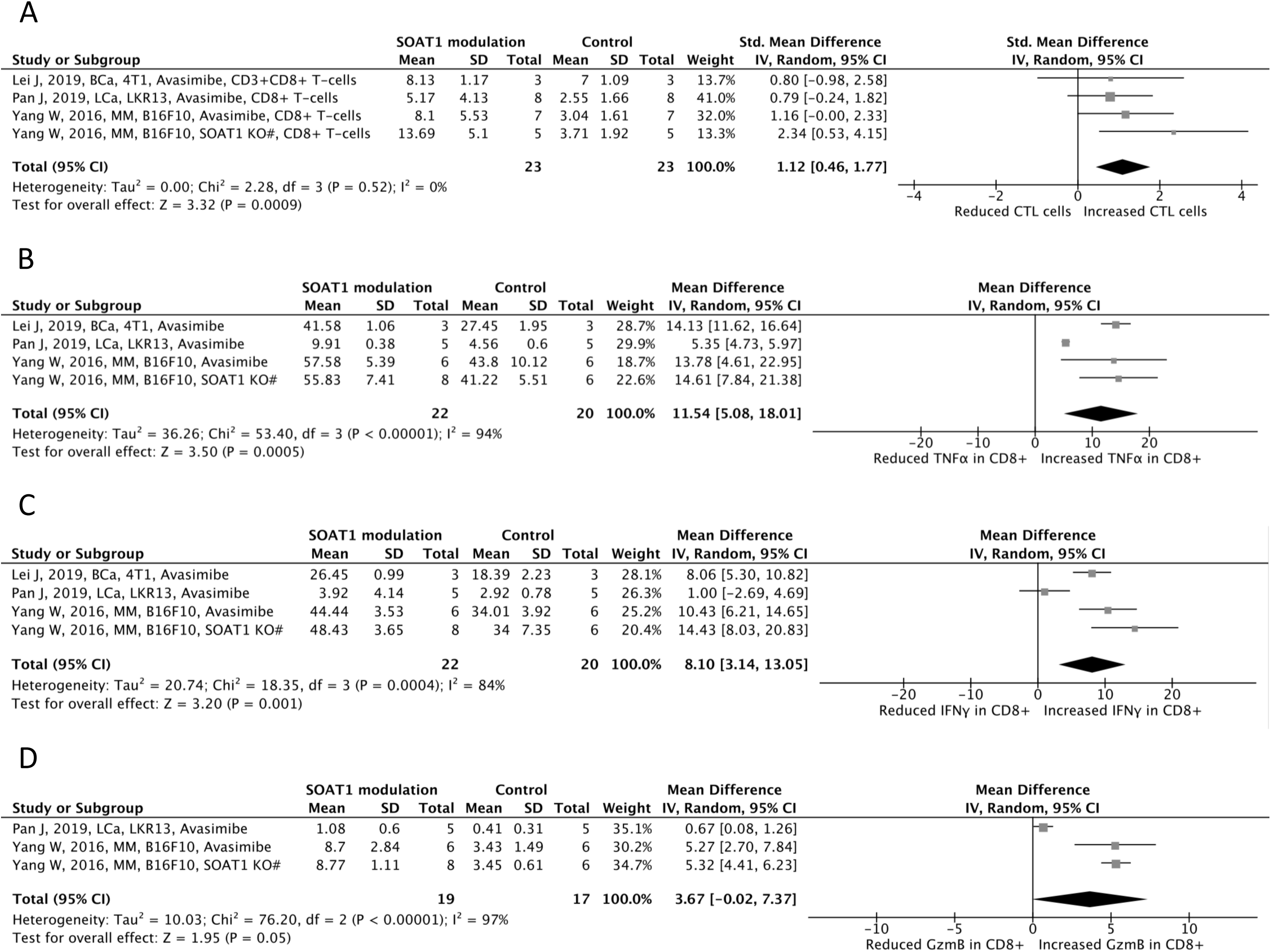
Forest plots of change in immune responses following disruption of SOAT. (A) Standardised mean difference between experimental and control in tumour infiltration of CD8+ cells. (B) Mean difference (percentage CD8+ cells) between experimental and control in TNFα expression in CD8+ cells. (C) Mean difference (percentage CD8+ cells) between experimental and control in IFNγ expression in CD8+ cells. (D) Mean difference (percentage CD8+ cells) between experimental and control in GzmB expression in CD8+ cells. # denotes modifications localized to T-cells.

Not only did SOAT1 disruption drive increased numbers of CTLs in some tumours, but CTLs had enhanced cytotoxic capabilities. We assessed differences in a range of cytotoxic effector cytokines across all appropriate studies and found without exception they were higher in tumours where SOAT had been inhibited: TNFα (MD = 11.54; 95% CI: 5.08 to 18.01; I^2^ = 94%; p = 0.0005; Fig6B), IFNγ (MD = 8.10; 95% CI: 3.14 to 13.05; I^2^ = 84%; p = 0.001 Fig6C), and cytotoxic effector molecule, GzmB (MD = 3.67; 95% CI: −0.02 to 7.37; I^2^ = 97%; p = 0.05 Fig6D).

#### 3.4.4 Activating invasion and metastasis

The ability of SOAT to drive metastatic colonisation was demonstrated in all four studies where metastasis was an endpoint (SMD = −2.21; 95% CI: −3.17 to −1.26; I^2^ = 57%; p < 0.00001; Fig7A). Avasimibe and avasimin supressed metastasis of breast cancer [75], pancreatic cancer [10], prostate cancer [66] and skin cancer [62] in the models. ShSOAT1 pre-treatment of MIA PaCa-2 pancreatic cancer xenografts reduced metastatic burden (lung metastasis x0.09, lymph metastasis x0.17) [10] and number of mice exhibiting metastatic lesions [54]. Furthermore, knockdown in MIA PaCa-2 xenograft cells reduced metastasis to the lung and lymph nodes. IV injection of LLC and B16F10 cells [62] paired with T-cell specific SOAT1 knockout reduced the metastatic potential of both cell lines (Table 4). Metastasis potential after avasimibe was also analysed by Hao et al. who found that a relatively small dose (2 mg/kg) was insufficient to reduce lung metastatic colonisation in lymphodepleted mice grafted IV with B16F10 cells [61]. Surprisingly, introduction of T-cells to this model lead to an increase in lung metastasis (Table 4). With the exception of this low dose study, SOAT activity in either tumour cells, T-cells, or both was associated with metastatic potential (Table 4).

**Figure 7.**
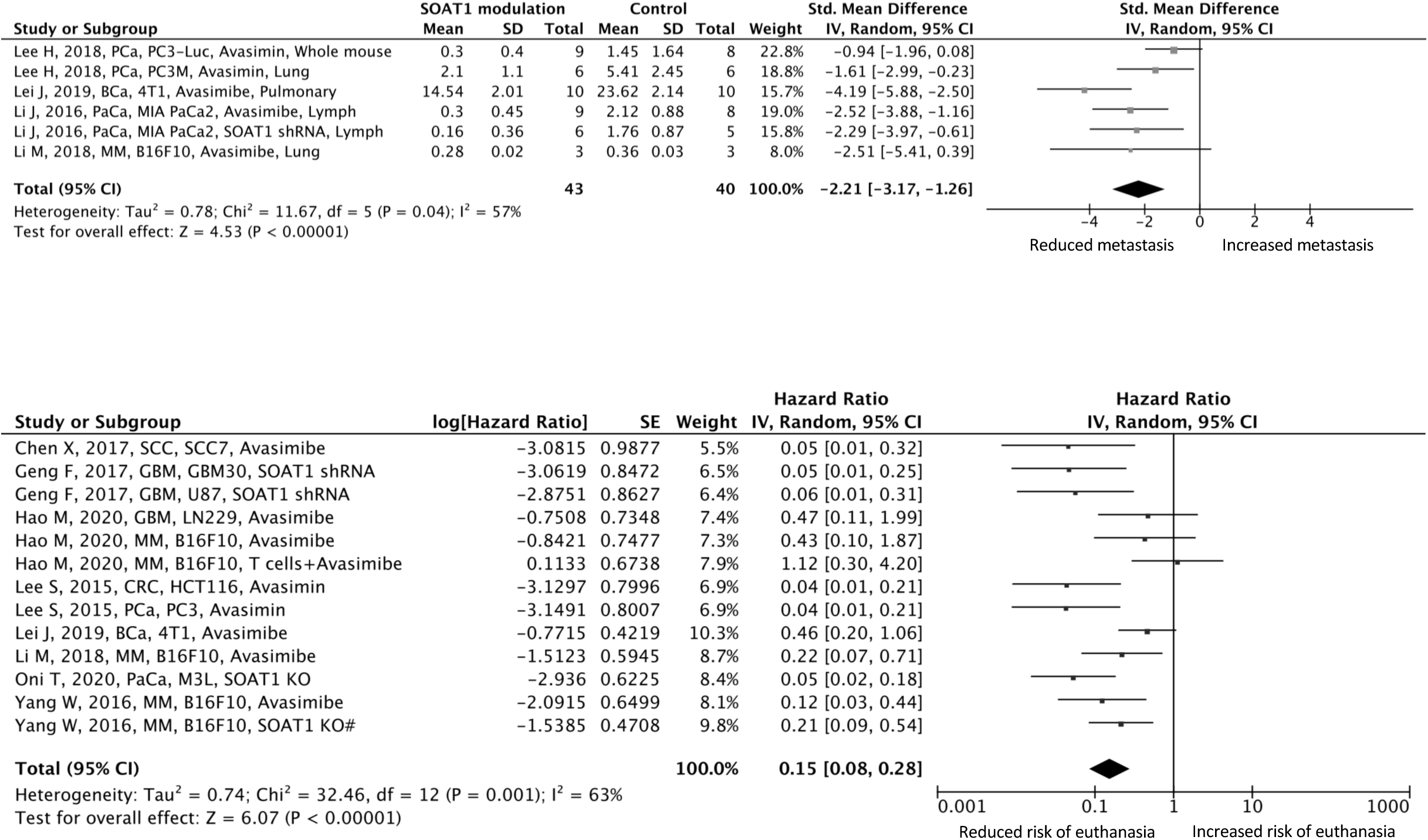
Forest plot showing changes in metastasis and risk of arrival at maximal tumour volume following disruption of SOAT. (A) Standardised mean difference between experimental and control number of metastases. (B) Differences shown as hazard ratios as calculated by Mantel-Haenszel between SOAT disruption test groups and control test groups. # denotes modifications localized to T-cells.

### 3.5 SOAT inhibition prolongs survival

Collectively, activation of cancer hallmarks increases tumour burden and is a prognostic indicator. Preclinical studies are bounded by ethical considerations that take into account animal suffering, which increases with tumour burden. Different regulatory agencies have different requirements on such experimental methods and typically state that when tumours reach a certain size, or animals lose a predetermined proportion of body weight, animals must be sacrificed. We utilised these data to calculate a novel hazard ratio function that describes the risk of the animal being euthanised based on local ethical requirements related to tumour burden. Thirteen experiments from seven studies [12, 39, 53, 54, 60–62, 75] provided data suitable for this analysis. Animals in intervention groups where SOAT1 function or expression was inhibited had an 85% reduction in risk of being euthanised earlier than the planned end of experimental period (HR = 0.15; 95% CI: 0.08 to 0.28; I^2^ = 63%; p < 0.00001; Fig7B).

### 3.6 Risk of bias analysis

#### 3.6.1 Study criteria

The quality of data included in our meta-analyses was measured using a multi-point survey that recorded data on transparency, scientific rigour, ethical animal research, and experimental reproducibility (Fig8). Every record was assessed by at least two independent researchers. Notably, fewer than half the studies validated that SOAT inhibition had been effective (Fig8A). The majority of the studies scoring poorly on this metric used avasimibe, which is well characterised. This perhaps also explains the lack of reporting on dosage rationale, with most studies administering avasimibe at a dose of 15mg/kg (66% of avasimibe treatments). Reporting on selection bias was lacking throughout all studies assessing SOAT disruption, with just 54% reporting randomisation of animals into test groups, only one study reporting assessor blinding to animal groups and none reporting randomised selection of animals for assessment. Furthermore, of the studies that reported randomisation of groups, none reported their method of randomisation. Additionally, no studies reported rationale behind the size of study groups, with this issue noted in previous meta-analyses on pre-clinical models of cancer [33, 76]. Comparably, studies assessing CE content in tissue exhibited poorer reporting on our risk of bias survey (Fig8B), however this is likely due to the papers within this cohort being considerably older (average publication date = 1986) than those assessing SOAT interventions in pre-clinical models (average publication date = 2018). Outside of these notable findings, reporting on other criteria was adequate and thus, risk of bias for study design was considered to be low. However, the chances of bias introduced through immunoblotting and immunohistochemistry is perhaps greater (Fig8C), with several studies lacking clarity of reporting on controls, statistical methods, and antibody validation.

**Figure 8.**
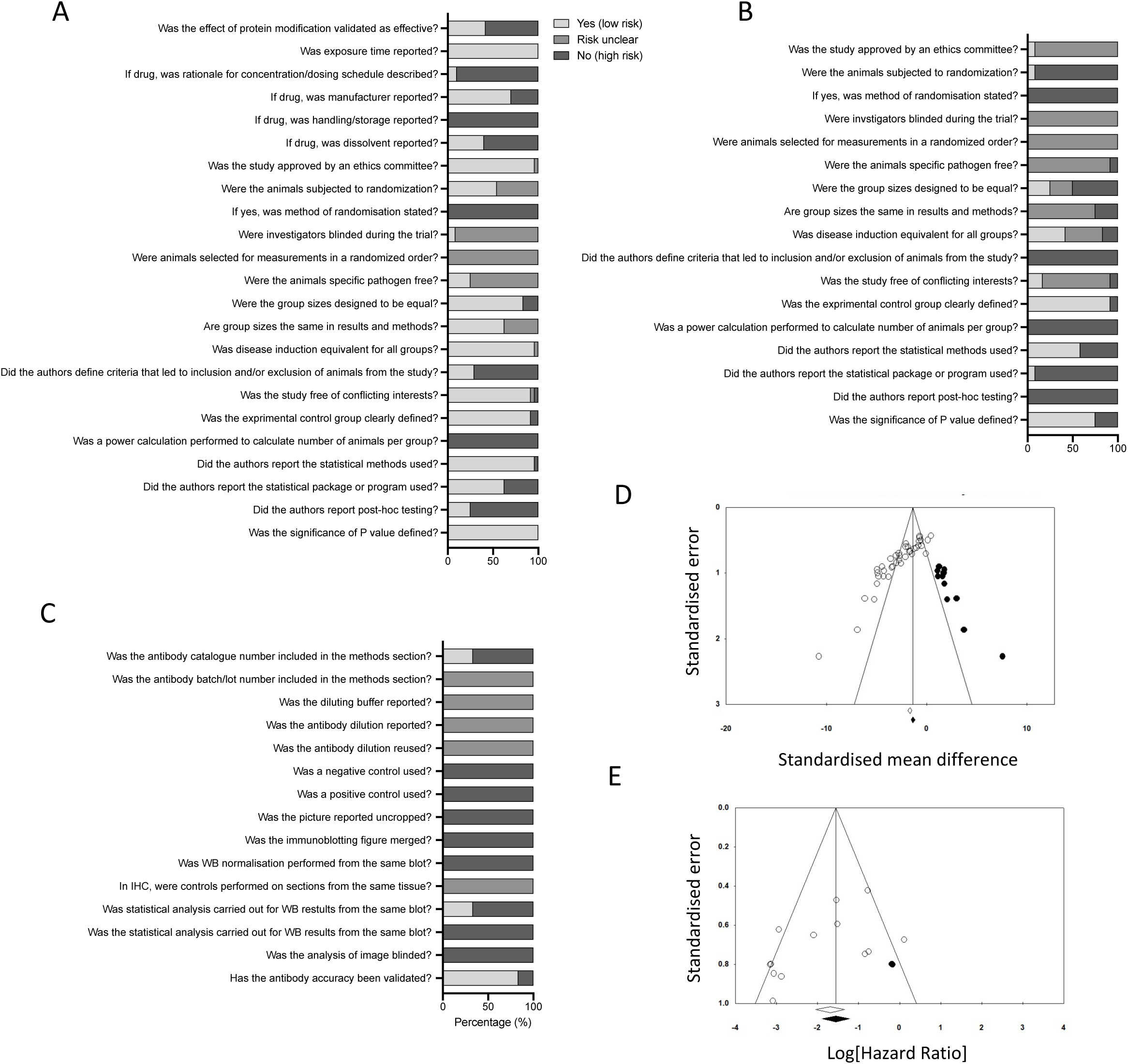
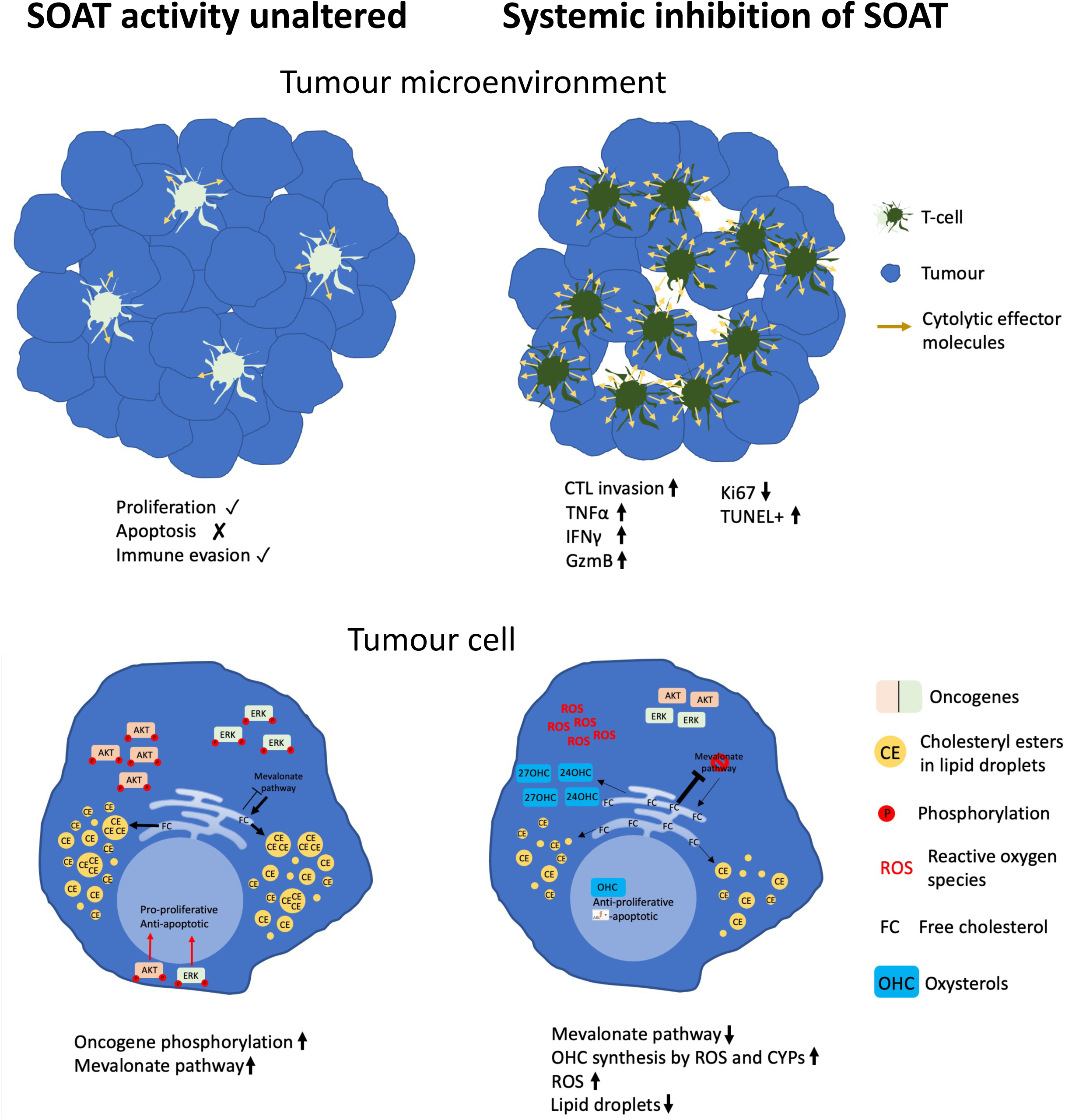
Risk of experimental and publication bias. (A) Adherence scores for animal research in studies assessing SOAT inhibition. (B) Adherence scores for animal research in studies measuring cholesterol ester content in tissue. (C) Adherence scores for immunoblotting in studies assessing SOAT inhibition. (D) Funnel plot to detect publication bias within SOAT1 tumour metrics dataset with trim and fill method applied to assess overestimation of SMD. (E) Funnel plot to detect publication bias within survival dataset with overestimation of hazard ratio determined through trim and fill analysis. Open dots indicate observed studies and closed dots indicate missing studies. Open diamond indicates observed change and the closed diamond indicates change after missing studies are factored in.

#### 3.6.2 Heterogeneity

There was high heterogeneity between cancer types (I^2^ = 82%) and within subgroup analysis. Some of this may be explained by differential expression of SOAT between cell types. For example, Jiang et al., used PDXs from six hepatocellular carcinomas, three with “high” and three with “low” SOAT1 expression. Avasimibe treatment led to significant impairment of tumour growth in the high, but not low, expressing tumours. This single study was the main contributor to the high heterogeneity observed in the liver cancer subgroup (I^2^ = 84%). However, SOAT expression is not the sole cause of the high heterogeneity. The prostate cancer studies exhibited high heterogeneity (I^2^ = 76%) despite all but one study examining the same cell line and originating from the same research group. Brain (I^2^ = 52%) and skin (I^2^ = 25%) cancers exhibit moderate to low levels of heterogeneity. When considering survival (section 3.5), despite the differences in cancer types and the range of methods utilised to modulate SOAT1 activity (drugs, RNAi, tumour to T-cell treatments) our analysis found only moderate heterogeneity (I^2^ = 63%).

#### 3.6.3 Publication Bias

Visual inspection of funnel plot for meta-analysis of tumour size and assessment of survival both suggested publication bias (Fig8D+E). This is likely driven by differences in methodological design between cancers. However, given that no individual cancer assessment was adequately powered (i.e., ≥ 10 studies) for an independent assessment of publication bias, a trim and fill method was used to estimate the degree of possible effect size overestimation across all cancers due to the suspected publication bias. Trim and fill method suggested the effect of SOAT inhibition on cancer size may be overestimated by 33% (Fig8D) while assessment of survival may be overestimated by 18% (Fig8E).

## 4. Discussion

This meta-analysis of 37 publications unequivocally shows that in animal cancer models CEs are elevated in cancer relative to normal tissue and inhibiting their synthesis reduces tumour burden. Importantly, in intervention groups tumours were smaller, less likely to metastasise, had reduced proliferative index, higher levels of apoptosis, and were more susceptible to destruction by cytotoxic T lymphocytes. These findings were highly significant and held true across all cancer sites evaluated including brain, liver, pancreas, prostate and skin.

Cholesterol plays a vital role in enabling efficient T-cell receptor clustering through its influence on membrane fluidity, leading to increased CTL activation. SOAT1 deficiency in CTLs leads to increased cholesterol content in the plasma membrane and enhanced T-cell receptor clustering, enhancing CTL cytotoxicity [61, 62]. Lei et al. proposed that enhanced cytotoxicity of CTLs is the primary driver of avasimibe’s anti-tumour effects [75]. Several lines of evidence support this. SOAT deficient T-cells exhibit enhanced cytotoxic potential against a variety of cancers *in vivo* and *in vitro* [55, 61, 62, 67, 75]. CD8+ cells pre-treated with avasimibe before B16F10 grafting led to higher levels of TNFα, IFNγ and GzmB [61] and complete eradication of tumour. Zhao et. al. found that siSOAT1 increased IFNγ expression in CAR-T cells and enhanced their cytotoxicity against MIA PaCa-2 xenografts [55]. *In vitro* cytotoxicity assays have also supported this hypothesis. SCC7 skin cancer cells are more susceptible to cytotoxic attack by CTLs if they are harvested from spleens of avasimibe exposed mice rather than from controls [60]. Moreover, CD8+ cells pre-treated with avasimibe and IV injected had grater cytotoxicity against B16F10 melanoma xenograft than mock pre-treated controls [61]. Furthermore, CTLs treated with SOAT1 specific inhibitor, K-604, induced greater EL-4 cell death than untreated CTLs [62]. This does not appear to be the case for all tumour types however, C26 colon cancer cells were considerably more resistant to CTL mediated cell death than B16F10 cells [53]. Pan et al. however, found that infiltrating T-cells exhibited no change in expression of cytotoxic markers or in ability to kill LKR13 cells after avasimibe exposure [67]. Interestingly, these cells express a KRAS mutant that is a key driver of T-cell immune checkpoint protein, PD-L1 [77], which drives T-cell exhaustion, and thus probably nullifies any effect from SOAT inhibition. Indeed, mutant KRAS inhibitors restored sensitivity to avasimibe and T-cell expression of TNFα, IFNγ and GzmB was increased in the dual treated cells [67].

CTLs are not the only anti-cancer mechanism likely to be at play. The xenograft data described here are gathered from nude mice that are broadly without T cells. A direct effect of SOAT inhibitors on tumour cells is also plausible. Cholesterol is not only esterified, but a range of enzymes can convert cholesterol into oxysterols by adding hydroxyl, keto, and epoxy moieties and avasimibe can generate reactive oxygen species [78] that also generate oxysterols. These oxysterols are anti-proliferative and pro-apoptotic in a range of cancer types [79]. SOAT1 and SOAT2 are both capable of esterifying oxysterols [1] and inhibition of SOAT2 leads to accumulation of 24-hydroxycholesterol and 26-hydroxycholesterol in Huh7 cells *in vitro* and *in vivo* [48]. Elevated oxysterol production within tumours may therefore explain the tumour suppressive effects of SOAT inhibition in the absence of a T-cell compartment. Interestingly, oxysterols also regulate T-cell function [80] so may act both directly on cancer cells and indirectly via the immune compartment. Consequently, these data suggest that SOAT may be acting to inhibit oxysterol’s anti-proliferative actions, and allowing cholesterol to be stored for use when needed and prevented from being converted into anti-proliferative oxysterols. However, the role of SOAT inhibition in regulating oxysterols was not considered in all but one [48] of the studies we found during our systematic searches. Measures of oxysterols should be considered vital in future work regarding SOAT inhibition in cancer.

Elevated CE concentration in cells appears also to influence cellular signalling cascades. For example, SOAT inhibition reduces phosphorylation of AKT [57–59] and ERK [40, 54, 59] oncogenes. Elevated intra-cellular free cholesterol resulting from SOAT inhibition was thought to be the cause of AKT dephosphorylation in pancreatic cells owing to downregulation of SREBP1 and LDL-receptor [58]. Reduced SREBP1 expression caused by SOAT1 inhibition has been reported in other pancreatic cell lines [59] and in glioblastoma cell lines [12]. Interestingly, addition of LXR synthetic ligand, T0901317, to pancreatic cancer cells inhibits phosphorylation of AKT [81], supporting the hypothesis that SOAT inhibition releases oxysterols. Oni et al. suggesting that SOAT1 mediated esterification of cholesterol prevents the negative feedback of the mevalonate pathway normally induced by free cholesterol. Loss of feedback prolongs cholesterol synthesis and other products of the pathway such as isoprenoids are produced. Isoprenoids themselves drive oncogenic activity of ERK [54], Ras and other GTP-binding proteins [82, 83].

Inhibition of SOAT activity may be a useful anti-cancer therapy, but several caveats are clear. Avasimibe performs poorly against tumours when given orally [39]. An IV route of administration for avasimibe or the more bioavailable avasimin may be more appropriate. Furthermore, avasimibe stimulates CYP3A4 activity in primary human hepatocytes [84] and given this detoxification enzyme is responsible for the metabolism of many chemotherapy agents, it is unsuitable as a combination therapy. Surprisingly, we found no evidence that induction of CYP3A11 (the mouse homologue of CYP3A4) was tested for in any of the 24 pre-clinical SOAT inhibition studies. This included five studies which examined and suggested SOAT inhibition should be performed alongside chemotherapy treatment [53, 57, 65, 67, 85]. SOAT inhibitors that do not activate CYP3A4 should be considered instead. For example, ATR-101 has no reported modulation of CYP3A4 activity and has already been investigated in adrenocortical carcinoma in a clinical trial. However, this trial found that the maximum safe dose of ATR-101 did not reduce cancer progression [29]. K-604 has been assessed for atherosclerosis treatment (NCT00851500) but results are currently unpublished.

Our risk of bias analysis identified a significant risk of publication bias in the field. Effect size was proportional to the number of animals in the study indicating that papers were more likely to be published if a significant effect had been found. Typically, in the absence of publication bias effect size is similar across studies, albeit with wider error margins in smaller studies. Correction by trim and fill method indicated that SOAT inhibition of tumour size and survival is probably overestimated by around a third and a fifth respectively. A caveat of this is that high heterogeneity between studies, which we observed (I^2^ = 71%), means the power to detect publication bias is reduced [86]. The trim and fill we performed may be skewed and the funnel plot asymmetry may result from inter-study differences rather than an under-reporting of either non-significant findings or studies with unexpected results [86]. We also found poor reporting of randomisation and assessor blinding, which increase the risk of bias [87]. Nevertheless, the effect we observed was strong, was found in many different types of measurement, across nearly all studies and cancer types, that SOAT inhibition is certainly linked to reduced tumour burden.

The role of cholesterol and cholesterol modifying interventions in cancer risk and progression has remained controversial for several years. The World Cancer Research Fund (WCRF) remain unable definitively to include or exclude cholesterol in the aetiology of cancer [88], even now, more than 20 years after the International Agency for Research on Cancer (IARC) indicated that evidence is inadequate [89]. Drugs and dietary factors that reduce cholesterol levels also reduce the risk of developing and/or dying from cancer in some situations [90–92]. Cholesterol is widely utilised, both structurally in the plasma membrane, and as a precursor for an array of hormones, steroids, and vitamins. The data we have explored and summarised here indicate that shifting the balance of cholesterol and manipulating its metabolism can have important consequences on cancer growth in animal models, which is at least in part mediated via the immune system. The summary we provide here indicates that the range of pharmacological inhibitors of cholesterol esterification that have not yet been evaluated in the cancer setting, pose attractive opportunities for drug-repurposing and chemoprevention of cancer.

## References

[1] S. Cases, S. Novak, Y.-W. Zheng, H.M. Myers, S.R. Lear, E. Sande, C.B. Welch, A.J. Lusis, T.A. Spencer, B.R. Krause, ACAT-2, a second mammalian acyl-CoA: cholesterol acyltransferase its cloning, expression, and characterization, Journal of Biological Chemistry 273(41) (1998) 26755–26764.

[2] X. Rousset, B. Vaisman, M. Amar, A.A. Sethi, A.T. Remaley, Lecithin: cholesterol acyltransferase: from biochemistry to role in cardiovascular disease, Current opinion in endocrinology, diabetes, and obesity 16(2) (2009) 163.

[3] T.-Y. Chang, C.C. Chang, S. Lin, C. Yu, B.-L. Li, A. Miyazaki, Roles of acyl-coenzyme A: cholesterol acyltransferase-1 and-2, Current opinion in lipidology 12(3) (2001) 289–296.

[4] S.E. Szedlacsek, E. Wasowicz, S.A. Hulea, H.I. Nishida, F.A. Kummerow, T. Nishida, Esterification of oxysterols by human plasma lecithin-cholesterol acyltransferase, Journal of Biological Chemistry 270(20) (1995) 11812–11819.

[5] A. Jonas, Lecithin cholesterol acyltransferase, Biochimica et Biophysica Acta (BBA)-Molecular and Cell Biology of Lipids 1529(1-3) (2000) 245–256.

[6] Y. Huang, Q. Jin, M. Su, F. Ji, N. Wang, C. Zhong, Y. Jiang, Y. Liu, Z. Zhang, J. Yang, Leptin promotes the migration and invasion of breast cancer cells by upregulating ACAT2, Cellular Oncology 40(6) (2017) 537–547.

[7] K. Matsumoto, Y. Fujiwara, R. Nagai, M. Yoshida, S. Ueda, Expression of two isozymes of acyl-coenzyme A: Cholesterol acyltransferase-1 and-2 in clear cell type renal cell carcinoma, International journal of urology 15(2) (2008) 166–170.

[8] Y. Jiang, A. Sun, Y. Zhao, W. Ying, H. Sun, X. Yang, B. Xing, W. Sun, L. Ren, B. Hu, C. Li, L. Zhang, G. Qin, M. Zhang, N. Chen, M. Zhang, Y. Huang, J. Zhou, Y. Zhao, M. Liu, X. Zhu, Y. Qiu, Y. Sun, C. Huang, M. Yan, M. Wang, W. Liu, F. Tian, H. Xu, J. Zhou, Z. Wu, T. Shi, W. Zhu, J. Qin, L. Xie, J. Fan, X. Qian, F. He, F. He, X. Qian, J. Qin, Y. Jiang, W. Ying, W. Sun, Y. Zhu, W. Zhu, Y. Wang, D. Yang, W. Liu, Q. Liu, X. Yang, B. Zhen, Z. Wu, J. Fan, H. Sun, J. Qian, T. Hong, L. Shen, B. Xing, P. Yang, H. Shen, L. Zhang, S. Cheng, J. Cai, X. Zhao, Y. Sun, T. Xiao, Y. Mao, X. Chen, D. Wu, L. Chen, J. Dong, H. Deng, M. Tan, Z. Wu, Q. Zhao, Z. Shen, X. Chen, Y. Gao, W. Sun, T. Wang, S. Liu, L. Lin, J. Zi, X. Lou, R. Zeng, Y. Wu, S. Cai, B. Jiang, A. Chen, Z. Li, F. Yang, X. Chen, Y. Sun, Q. Wang, Y. Zhang, G. Wang, Z. Chen, W. Qin, Z. Li, C. Chinese Human Proteome Project, Proteomics identifies new therapeutic targets of early-stage hepatocellular carcinoma, Nature 567(7747) (2019) 257-261.

[9] K.C. Chi, W.C. Tsai, C.L. Wu, T.Y. Lin, D.Y. Hueng, An Adult Drosophila Glioma Model for Studying Pathometabolic Pathways of Gliomagenesis, Mol Neurobiol 56(6) (2019) 4589–4599.

[10] J. Li, D. Gu, S.S.Y. Lee, B. Song, S. Bandyopadhyay, S. Chen, S.F. Konieczny, T.L. Ratliff, X. Liu, J. Xie, J.X. Cheng, Abrogating cholesterol esterification suppresses growth and metastasis of pancreatic cancer Oncogene 35(50) (2016) 6378–6388.

[11] A.M.F. Lacombe, I.C. Soares, B.M.d.P. Mariani, M.Y. Nishi, J.E. Bezerra-Neto, H.d.S. Charchar, V.B. Brondani, F. Tanno, V. Srougi, J.L. Chambo, R.M. Costa de Freitas, B.B. Mendonca, A.O. Hoff, M.Q. Almeida, I. Weigand, M. Kroiss, M.C.N. Zerbini, M.C.B.V. Fragoso, Sterol O-Acyl Transferase 1 as a Prognostic Marker of Adrenocortical Carcinoma, Cancers 12(1) (2020) 247.

[12] F. Geng, X. Cheng, X. Wu, J.Y. Yoo, C. Cheng, J.Y. Guo, X. Mo, P. Ru, B. Hurwitz, S.-H. Kim, Inhibition of SOAT1 suppresses glioblastoma growth via blocking SREBP-1–mediated lipogenesis, Clinical Cancer Research 22(21) (2016) 5337–5348.

[13] J. Li, S. Ren, H.-l. Piao, F. Wang, P. Yin, C. Xu, X. Lu, G. Ye, Y. Shao, M. Yan, Integration of lipidomics and transcriptomics unravels aberrant lipid metabolism and defines cholesteryl oleate as potential biomarker of prostate cancer, Scientific reports 6(1) (2016) 1–11.

[14] J. Long, P. Chen, J. Lin, Y. Bai, X. Yang, J. Bian, Y. Lin, D. Wang, X. Yang, Y. Zheng, DNA methylation-driven genes for constructing diagnostic, prognostic, and recurrence models for hepatocellular carcinoma, Theranostics 9(24) (2019) 7251.

[15] G. Ouyang, B. Yi, G. Pan, X. Chen, A robust twelve-gene signature for prognosis prediction of hepatocellular carcinoma, Cancer Cell International 20 (2020) 1–18.

[16] S.P. Pattanayak, P. Sunita, P.M. Mazumder, Restorative effect of Dendrophthoe falcata (Lf) Ettingsh on lipids, lipoproteins, and lipid-metabolizing enzymes in DMBA-induced mammary gland carcinogenesis in Wistar female rats, Comparative Clinical Pathology 23(4) (2014) 1013–1022.

[17] C. Thirunavukkarasu, K. Selvedhiran, J. Prince Vijaya Singh, P. Senthilnathan, D. Sakthisekaran, Effect of sodium selenite on lipids and lipid-metabolizing enzymes in N-nitrosodiethylamine-induced hepatoma-bearing rats, The Journal of Trace Elements in Experimental Medicine: The Official Publication of the International Society for Trace Element Research in Humans 16(1) (2003) 1–15.

[18] K. Veena, P. Shanthi, P. Sachdanandam, The biochemical alterations following administration of Kalpaamruthaa and Semecarpus anacardium in mammary carcinoma, Chemico-biological interactions 161(1) (2006) 69–78.

[19] M.R. Paillasse, P. de Medina, G. Amouroux, L. Mhamdi, M. Poirot, S. Silvente-Poirot, Signaling through cholesterol esterification: a new pathway for the cholecystokinin 2 receptor involved in cell growth and invasion, Journal of lipid research 50(11) (2009) 2203–2211.

[20] P. de Medina, S. Genovese, M.R. Paillasse, M. Mazaheri, S. Caze-Subra, K. Bystricky, M. Curini, S. Silvente-Poirot, F. Epifano, M. Poirot, Auraptene is an inhibitor of cholesterol esterification and a modulator of estrogen receptors, Molecular pharmacology 78(5) (2010) 827–836.

[21] F. Khallouki, R.W. Owen, S. Silvente-Poirot, M. Poirot, Bryonolic acid blocks cancer cell clonogenicity and invasiveness through the inhibition of fatty acid: cholesteryl ester formation, Biomedicines 6(1) (2018) 21.

[22] H.T. Lee, D.R. Sliskovic, J.A. Picard, B.D. Roth, W. Wierenga, J.L. Hicks, R.F. Bousley, K.L. Hamelehle, R. Homan, C. Speyer, Inhibitors of acyl-CoA: cholesterol O-acyl transferase (ACAT) as hypocholesterolemic agents. CI-1011: an acyl sulfamate with unique cholesterol-lowering activity in animals fed noncholesterol-supplemented diets, Journal of medicinal chemistry 39(26) (1996) 5031-5034.

[23] N. Terasaka, A. Miyazaki, N. Kasanuki, K. Ito, N. Ubukata, T. Koieyama, K. Kitayama, T. Tanimoto, N. Maeda, T. Inaba, ACAT inhibitor pactimibe sulfate (CS-505) reduces and stabilizes atherosclerotic lesions by cholesterol-lowering and direct effects in apolipoprotein E-deficient mice, Atherosclerosis 190(2) (2007) 239–247.

[24] M. Ikenoya, Y. Yoshinaka, H. Kobayashi, K. Kawamine, K. Shibuya, F. Sato, K. Sawanobori, T. Watanabe, A. Miyazaki, A selective ACAT-1 inhibitor, K-604, suppresses fatty streak lesions in fat-fed hamsters without affecting plasma cholesterol levels, Atherosclerosis 191(2) (2007) 290-297.

[25] J.-C. Tardif, J. Grégoire, P.L. L’Allier, T.J. Anderson, O. Bertrand, F. Reeves, L.M. Title, F. Alfonso, E. Schampaert, A. Hassan, Effects of the acyl coenzyme A: cholesterol acyltransferase inhibitor avasimibe on human atherosclerotic lesions, Circulation 110(21) (2004) 3372–3377.

[26] F.J. Raal, A.D. Marais, E. Klepack, J. Lovalvo, R. McLain, T. Heinonen, Avasimibe, an ACAT inhibitor, enhances the lipid lowering effect of atorvastatin in subjects with homozygous familial hypercholesterolemia, Atherosclerosis 171(2) (2003) 273–279.

[27] H.J. Lee, S.H. Yue, J.J. Li, S.Y. Lee, T. Shao, B. Song, L. Cheng, T.A. Masterson, X.Q. Liu, T.L. Ratliff, J.X. Cheng, Cholesteryl ester accumulation induced by PTEN loss and PI3K/AKT activation underlies human prostate cancer aggressiveness Molecular Cancer Therapeutics 14(7) (2015).

[28] C.R. LaPensee, J.E. Mann, W.E. Rainey, V. Crudo, S.W. Hunt III, G.D. Hammer, ATR-101, a selective and potent inhibitor of acyl-CoA acyltransferase 1, induces apoptosis in H295R adrenocortical cells and in the adrenal cortex of dogs, Endocrinology 157(5) (2016) 1775–1788.

[29] D.C. Smith, M. Kroiss, E. Kebebew, M.A. Habra, R. Chugh, B.J. Schneider, M. Fassnacht, P. Jafarinasabian, M.M. Ijzerman, V.H. Lin, A phase 1 study of nevanimibe HCl, a novel adrenal-specific sterol O-acyltransferase 1 (SOAT1) inhibitor, in adrenocortical carcinoma, Investigational new drugs 38(5) (2020) 1421–1429.

[30] K. Kitayama, T. Tanimoto, T. Koga, N. Terasaka, T. Fujioka, T. Inaba, Importance of acyl-coenzyme A: cholesterol acyltransferase 1/2 dual inhibition for anti-atherosclerotic potency of pactimibe, European journal of pharmacology 540(1-3) (2006) 121–130.

[31] M.C. Meuwese, E. de Groot, R. Duivenvoorden, M.D. Trip, L. Ose, F.J. Maritz, D.C. Basart, J.J. Kastelein, R. Habib, M.H. Davidson, A.H. Zwinderman, L.R. Schwocho, E.A. Stein, C. Investigators, ACAT inhibition and progression of carotid atherosclerosis in patients with familial hypercholesterolemia: the CAPTIVATE randomized trial, JAMA 301(11) (2009) 1131–9.

[32] D. Jackson, R. Turner, Power analysis for random-effects meta-analysis, Research synthesis methods 8(3) (2017) 290–302.

[33] G. Cioccoloni, C. Soteriou, A. Websdale, L. Wallis, M.A. Zulyniak, J.L. Thorne, Phytosterols and phytostanols and the hallmarks of cancer in model organisms: A systematic review and meta-analysis, Critical Reviews in Food Science and Nutrition (2020) 1–21.

[34] S.P. Alexander, R.E. Roberts, B.R. Broughton, C.G. Sobey, C.H. George, S.C. Stanford, G. Cirino, J.R. Docherty, M.A. Giembycz, D. Hoyer, Goals and practicalities of immunoblotting and immunohistochemistry: A guide for submission to the British Journal of Pharmacology, Wiley Online Library, 2018.

[35] BJP, Declaration of transparency and scientific rigour: checklist for animal experimentation, Br J Pharmacol 175(13) (2018) 2711.

[36] C.R. Hooijmans, M.M. Rovers, R.B.M. de Vries, M. Leenaars, M. Ritskes-Hoitinga, M.W. Langendam, SYRCLE’s risk of bias tool for animal studies, BMC Medical Research Methodology 14(1) (2014) 43.

[37] A.C. Ross, K. Go, J. Heider, G. Rothblat, Selective inhibition of acyl coenzyme A: cholesterol acyltransferase by compound 58-035, Journal of Biological Chemistry 259(2) (1984) 815–819.

[38] M. Weng, H. Zhang, W. Hou, Z. Sun, J. Zhong, C. Miao, ACAT2 Promotes Cell Proliferation and Associates with Malignant Progression in Colorectal Cancer, OncoTargets and therapy 13 (2020) 3477.

[39] S.S.Y. Lee, J.J. Li, J.N. Tai, T.L. Ratliff, K. Park, J.X. Cheng, Avasimibe Encapsulated in Human Serum Albumin Blocks Cholesterol Esterification for Selective Cancer Treatment, Acs Nano 9(3) (2015) 2420–2432.

[40] L. Wang, Y. Liu, G. Yu, Avasimibe inhibits tumor growth by targeting foxm1-akr1c1 in osteosarcoma, OncoTargets and therapy 12 (2019) 815.

[41] G.P.H. van Heushen, T.P. van der Krift, K.Y. Hostetler, K.W. Wirtz, Effect of nonspecific phospholipid transfer protein on cholesterol esterification in microsomes from Morris hepatomas, Cancer research 43(9) (1983) 4207–4210.

[42] D. Harry, H. Morris, N. McINTYRE, Cholesterol biosynthesis in transplantable hepatomas: evidence for impairment of uptake and storage of dietary cholesterol, Journal of lipid research 12(3) (1971) 313–317.

[43] J.Y. Liu, W.Q. Fu, X.J. Zheng, W. Li, L.W. Ren, J.H. Wang, C. Yang, G.H. Du, Avasimibe exerts anticancer effects on human glioblastoma cells via inducing cell apoptosis and cell cycle arrest, Acta Pharmacologica Sinica (2020).

[44] Y. Luo, L. Liu, X. Li, Y. Shi, Avasimibe inhibits the proliferation, migration and invasion of glioma cells by suppressing linc00339, Biomedicine & Pharmacotherapy 130 (2020) 110508.

[45] Y.H. Cheng, R.E. Kerppola, T.K. Kerppola, ATR-101 disrupts mitochondrial functions in adrenocortical carcinoma cells and in vivo, Endocrine-Related Cancer 23(4) (2016) 1–19.

[46] V.C. Prabhu, Glioblastoma Multiforme, 16/05/2021 https://www.aans.org/en/Patients/Neurosurgical-Conditions-and-Treatments/Glioblastoma-Multiforme#:~:text=Prevalence%20and%20Incidence,men%20as%20compared%20to%20women. (2021).

[47] M. Ayala-Ramirez, S. Jasim, L. Feng, S. Ejaz, F. Deniz, N. Busaidy, S.G. Waguespack, A. Naing, K. Sircar, C.G. Wood, Adrenocortical carcinoma: clinical outcomes and prognosis of 330 patients at a tertiary care center, European journal of endocrinology/European Federation of Endocrine Societies 169(6) (2013) 891.

[48] M. Lu, X.H. Hu, Q. Li, Y. Xiong, G.J. Hu, J.J. Xu, X.N. Zhao, X.X. Wei, C.C. Chang, Y.K. Liu, F.J. Nan, J. Li, T.Y. Chang, B.L. Song, B.L. Li, A specific cholesterol metabolic pathway is established in a subset of HCCs for tumor growth, J Mol Cell Biol 5(6) (2013) 404–15.

[49] H. Sung, J. Ferlay, R.L. Siegel, M. Laversanne, I. Soerjomataram, A. Jemal, F. Bray, Global cancer statistics 2020: GLOBOCAN estimates of incidence and mortality worldwide for 36 cancers in 185 countries, CA Cancer J Clin (2021).

[50] M. Uhlen, C. Zhang, S. Lee, E. Sjöstedt, L. Fagerberg, G. Bidkhori, R. Benfeitas, M. Arif, Z. Liu, F. Edfors, A pathology atlas of the human cancer transcriptome, Science 357(6352: https://www.proteinatlas.org/ENSG00000057252-SOAT1/cell) (2017).

[51] M. Uhlen, C. Zhang, S. Lee, E. Sjöstedt, L. Fagerberg, G. Bidkhori, R. Benfeitas, M. Arif, Z. Liu, F. Edfors, A pathology atlas of the human cancer transcriptome, Science 357(6352: https://www.proteinatlas.org/ENSG00000167780-SOAT2/cell) (2017).

[52] B.-L. Song, C.-H. Wang, X.-M. Yao, L. Yang, W.-J. Zhang, Z.-Z. Wang, X.-N. Zhao, J.-B. Yang, W. Qi, X.-Y. Yang, Human acyl-CoA: cholesterol acyltransferase 2 gene expression in intestinal Caco-2 cells and in hepatocellular carcinoma, Biochemical Journal 394(3) (2006) 617–626.

[53] M. Li, Y.T. Yang, J.J. Wei, X.L. Cun, Z.Z. Lu, Y. Qiu, Z.R. Zhang, Q. He, Enhanced chemo-immunotherapy against melanoma by inhibition of cholesterol esterification in CD8(+) T cells, Nanomedicine-Nanotechnology Biology and Medicine 14(8) (2018) 2541–2550.

[54] T.E. Oni, G. Biffi, L.A. Baker, Y. Hao, C. Tonelli, T.D. Somerville, A. Deschênes, P. Belleau, C.-i. Hwang, F.J. Sánchez-Rivera, SOAT1 promotes mevalonate pathway dependency in pancreatic cancer, Journal of Experimental Medicine 217(9) (2020).

[55] L. Zhao, Y. Liu, F. Zhao, Y. Jin, J. Feng, R. Geng, J. Sun, L. Kang, L. Yu, Y. Wei, Inhibition of cholesterol esterification enzyme enhances the potency of human chimeric antigen receptor T cells against pancreatic carcinoma, Molecular Therapy-Oncolytics 16 (2020) 262–271.

[56] C.R. UK, Survival. Available from: https://www.cancerresearchuk.org/about-cancer/pancreatic-cancer/survival [Accessed 6th August 2020].

[57] J. Li, X. Qu, J. Tian, J.T. Zhang, J.X. Cheng, Cholesterol esterification inhibition and gemcitabine synergistically suppress pancreatic ductal adenocarcinoma proliferation, PLoS ONE 13(2) (2018).

[58] S.H. Yue, J.J. Li, S.Y. Lee, H.J. Lee, T. Shao, B. Song, L. Cheng, T.A. Masterson, X.Q. Liu, T.L. Ratliff, J.X. Cheng, Cholesteryl Ester Accumulation Induced by PTEN Loss and PI3K/AKT Activation Underlies Human Prostate Cancer Aggressiveness, Cell Metabolism 19(3) (2014) 393–406.

[59] Y. Liu, Y. Wang, S. Hao, Y. Qin, Y. Wu, Knockdown of sterol O-acyltransferase 1 (SOAT1) suppresses SCD1-mediated lipogenesis and cancer procession in prostate cancer, Prostaglandins & Other Lipid Mediators 153 (2021) 106537.

[60] X. Chen, Q. Song, L. Xia, X. Xu, Synergy of dendritic cell vaccines and avasimibe in treatment of head and neck cancer in mice, Medical Science Monitor 23 (2017) 4471–4476.

[61] M. Hao, S. Hou, W. Li, K. Li, L. Xue, Q. Hu, L. Zhu, Y. Chen, H. Sun, C. Ju, Combination of metabolic intervention and T cell therapy enhances solid tumor immunotherapy, Science Translational Medicine 12(571) (2020).

[62] W. Yang, Y. Bai, Y. Xiong, J. Zhang, S. Chen, X. Zheng, X. Meng, L. Li, J. Wang, C. Xu, C. Yan, L. Wang, C.C. Chang, T.Y. Chang, T. Zhang, P. Zhou, B.L. Song, W. Liu, S.C. Sun, X. Liu, B.L. Li, C. Xu, Potentiating the antitumour response of CD8(+) T cells by modulating cholesterol metabolism, Nature 531(7596) (2016) 651-5.

[63] C.L. Sawyers, Chronic myeloid leukemia, New England Journal of Medicine 340(17) (1999) 1330–1340.

[64] D. Cilloni, G. Saglio, Molecular pathways: Bcr-abl, Clinical Cancer Research 18(4) (2012) 930–937.

[65] S. Bandyopadhyay, J. Li, E. Traer, J.W. Tyner, A. Zhou, S.T. Oh, J.X. Cheng, Cholesterol esterification inhibition and imatinib treatment synergistically inhibit growth of BCR-ABL mutation-independent resistant chronic myelogenous leukemia, PLoS One 12(7) (2017) e0179558.

[66] H.J. Lee, J. Li, R.E. Vickman, J.J. Li, R. Liu, A.C. Durkes, B.D. Elzey, S.H. Yue, X.Q. Liu, T.L. Ratliff, J.X. Cheng, Cholesterol Esterification Inhibition Suppresses Prostate Cancer Metastasis by Impairing the Wnt/beta-catenin Pathway Molecular Cancer Research 16(6) (2018) 974–985.

[67] J. Pan, Q. Zhang, K. Palen, L. Wang, L. Qiao, B. Johnson, S. Sei, R.H. Shoemaker, R.A. Lubet, Y. Wang, M. You, Potentiation of Kras peptide cancer vaccine by avasimibe, a cholesterol modulator, EBioMedicine 49 (2019) 72–81.

[68] R. Menegaz, M. Michelin, R. Etchebehere, P. Fernandes, E. Murta, Peri-and intratumoral T and B lymphocytic infiltration in breast cancer, European journal of gynaecological oncology 29(4) (2008) 321.

[69] J. Galon, A. Costes, F. Sanchez-Cabo, A. Kirilovsky, B. Mlecnik, C. Lagorce-Pagès, M. Tosolini, M. Camus, A. Berger, P. Wind, Type, density, and location of immune cells within human colorectal tumors predict clinical outcome, Science 313(5795) (2006) 1960-1964.

[70] O. Kawai, G. Ishii, K. Kubota, Y. Murata, Y. Naito, T. Mizuno, K. Aokage, N. Saijo, Y. Nishiwaki, A. Gemma, Predominant infiltration of macrophages and CD8+ T cells in cancer nests is a significant predictor of survival in stage IV nonsmall cell lung cancer, Cancer: Interdisciplinary International Journal of the American Cancer Society 113(6) (2008) 1387–1395.

[71] C.G. Clemente, M.C. Mihm Jr, R. Bufalino, S. Zurrida, P. Collini, N. Cascinelli, Prognostic value of tumor infiltrating lymphocytes in the vertical growth phase of primary cutaneous melanoma, Cancer: Interdisciplinary International Journal of the American Cancer Society 77(7) (1996) 1303–1310.

[72] E. Richardsen, R. Uglehus, J. Due, C. Busch, L.T. Busund, The prognostic impact of M-CSF, CSF-1 receptor, CD68 and CD3 in prostatic carcinoma, Histopathology 53(1) (2008) 30-38.

[73] P.P. Lee, C. Yee, P.A. Savage, L. Fong, D. Brockstedt, J.S. Weber, D. Johnson, S. Swetter, J. Thompson, P.D. Greenberg, M. Roederer, M.M. Davis, Characterization of circulating T cells specific for tumor-associated antigens in melanoma patients, Nature Medicine 5(6) (1999) 677–685.

[74] M.R. Dowling, A. Kan, S. Heinzel, J.M. Marchingo, P.D. Hodgkin, E.D. Hawkins, Regulatory T cells suppress effector T cell proliferation by limiting division destiny, Frontiers in immunology 9 (2018) 2461.

[75] J. Lei, H.J. Wang, D.M. Zhu, Y.B. Wan, L. Yin, Combined effects of avasimibe immunotherapy, doxorubicin chemotherapy, and metal-organic frameworks nanoparticles on breast cancer, Journal of Cellular Physiology 235(5) (2020) 4814–4823.

[76] J. Mattina, N. MacKinnon, V.C. Henderson, D. Fergusson, J. Kimmelman, Design and reporting of targeted anticancer preclinical studies: a meta-analysis of animal studies investigating sorafenib antitumor efficacy, Cancer research 76(16) (2016) 4627–4636.

[77] H. Sumimoto, A. Takano, K. Teramoto, Y. Daigo, RAS–mitogen-activated protein kinase signal is required for enhanced PD-L1 expression in human lung cancers, PloS one 11(11) (2016) e0166626.

[78] V.N. Ayyagari, X. Wang, P.L. Diaz-Sylvester, K. Groesch, L. Brard, Assessment of acyl-CoA cholesterol acyltransferase (ACAT-1) role in ovarian cancer progression—An in vitro study, Plos one 15(1) (2020) e0228024.

[79] C.-P. Chuu, H.-P. Lin, Antiproliferative effect of LXR agonists T0901317 and 22 (R)-hydroxycholesterol on multiple human cancer cell lines, Anticancer research 30(9) (2010) 3643–3648.

[80] A.E. Baek, Y.-R.A. Yu, S. He, S.E. Wardell, C.-Y. Chang, S. Kwon, R.V. Pillai, H.B. McDowell, J.W. Thompson, L.G. Dubois, P.M. Sullivan, J.K. Kemper, M.D. Gunn, D.P. McDonnell, E.R. Nelson, The cholesterol metabolite 27 hydroxycholesterol facilitates breast cancer metastasis through its actions on immune cells, Nature Communications 8(1) (2017) 864.

[81] A. Pommier, G. Alves, E. Viennois, S. Bernard, Y. Communal, B. Sion, G. Marceau, C. Damon, K. Mouzat, F. Caira, Liver X Receptor activation downregulates AKT survival signaling in lipid rafts and induces apoptosis of prostate cancer cells, Oncogene 29(18) (2010) 2712–2723.

[82] A. Ridley, RhoA, RhoB and RhoC have different roles in cancer cell migration, Journal of microscopy 251(3) (2013) 242–249.

[83] R. Riscal, N. Skuli, M.C. Simon, Even cancer cells watch their cholesterol!, Molecular cell 76(2) (2019) 220–231.

[84] J. Sahi, M.A. Milad, X. Zheng, K.A. Rose, H. Wang, L. Stilgenbauer, D. Gilbert, S. Jolley, R.H. Stern, E.L. LeCluyse, Avasimibe induces CYP3A4 and multiple drug resistance protein 1 gene expression through activation of the pregnane X receptor, Journal of Pharmacology and Experimental Therapeutics 306(3) (2003) 1027–1034.

[85] M. Bi, X. Qiao, H. Zhang, H. Wu, Z. Gao, H. Zhou, M. Shi, Y. Wang, J. Yang, J. Hu, W. Liang, Y. Liu, X. Qiao, S. Zhang, Z. Zhao, Effect of inhibiting ACAT-1 expression on the growth and metastasis of Lewis lung carcinoma, Oncol Lett 18(2) (2019) 1548–1556.

[86] J. Lau, J.P. Ioannidis, N. Terrin, C.H. Schmid, I. Olkin, The case of the misleading funnel plot, Bmj 333(7568) (2006) 597-600.

[87] J.A. Hirst, J. Howick, J.K. Aronson, N. Roberts, R. Perera, C. Koshiaris, C. Heneghan, The need for randomization in animal trials: an overview of systematic reviews, PLoS One 9(6) (2014) e98856.

[88] WCRF/AICR, Diet, Nutrition, Physical Activity and Cancer: a Global Perspective. Continuous Update Project Expert Report., World Cancer Research Fund/American Institute for Cancer Research, 2018.

[89] IARC, Cholesterol, IARC, publications.iarc.fr, 1996.

[90] G. Cioccoloni, C. Soteriou, A. Websdale, L. Wallis, M.A. Zulyniak, J.L. Thorne, Phytosterols and phytostanols and the hallmarks of cancer in model organisms: A systematic review and meta-analysis, Crit Rev Food Sci Nutr (2020) 1–21.

[91] L. Jiang, X. Zhao, J. Xu, C. Li, Y. Yu, W. Wang, L. Zhu, The Protective Effect of Dietary Phytosterols on Cancer Risk: A Systematic Meta-Analysis, Journal of Oncology 2019 (2019) 11.

[92] B. Liu, Z. Yi, X. Guan, Y.X. Zeng, F. Ma, The relationship between statins and breast cancer prognosis varies by statin type and exposure time: a meta-analysis, Breast Cancer Res Treat 164(1) (2017) 1–11.

[93] W. Insull Jr, M. Koren, J. Davignon, D. Sprecher, H. Schrott, L.M. Keilson, A.S. Brown, C.A. Dujovne, M.H. Davidson, R. McLain, Efficacy and short-term safety of a new ACAT inhibitor, avasimibe, on lipids, lipoproteins, and apolipoproteins, in patients with combined hyperlipidemia, Atherosclerosis 157(1) (2001) 137–144.

[94] D.C. Smith, M. Kroiss, E. Kebebew, M.A. Habra, R. Chugh, B.J. Schneider, M. Fassnacht, P. Jafarinasabian, M.M. Ijzerman, V.H. Lin, P. Mohideen, A. Naing, A phase 1 study of nevanimibe HCl, a novel adrenal-specific sterol O-acyltransferase 1 (SOAT1) inhibitor, in adrenocortical carcinoma, Investigational New Drugs 38(5) (2020) 1421–1429.

[95] D. El-Maouche, D.P. Merke, M.G. Vogiatzi, A.Y. Chang, A.F. Turcu, E.G. Joyal, V.H. Lin, L. Weintraub, M.R. Plaunt, P. Mohideen, R.J. Auchus, A Phase 2, Multicenter Study of Nevanimibe for the Treatment of Congenital Adrenal Hyperplasia, The Journal of Clinical Endocrinology & Metabolism 105(8) (2020) 2771–2778.

[96] M.C. Meuwese, E. de Groot, R. Duivenvoorden, M.D. Trip, L. Ose, F.J. Maritz, D.C.G. Basart, J.J.P. Kastelein, R. Habib, M.H. Davidson, A.H. Zwinderman, L.R. Schwocho, E.A. Stein, f.t. Captivate Investigators, ACAT Inhibition and Progression of Carotid Atherosclerosis in Patients With Familial Hypercholesterolemia: The CAPTIVATE Randomized Trial, JAMA 301(11) (2009) 1131–1139.

[97] T. Ohshiro, L.L. Rudel, S. Ōmura, H. Tomoda, Selectivity of microbial acyl-CoA: cholesterol acyltransferase inhibitors toward isozymes, The Journal of antibiotics 60(1) (2007) 43–51.

[98] A.T. Lada, M. Davis, C. Kent, J. Chapman, H. Tomoda, S. Omura, L.L. Rudel, Identification of ACAT1-and ACAT2-specific inhibitors using a novel, cell-based fluorescence assay: individual ACAT uniqueness, Journal of lipid research 45(2) (2004) 378–386.

[99] I. Tabas, L.L. Chen, J.W. Clader, A.T. McPhail, D.A. Burnett, P. Bartner, P.R. Das, B.N. Pramanik, M.S. Puar, S.J. Feinmark, Rabbit and human liver contain a novel pentacyclic triterpene ester with acyl-CoA: cholesterol acyltransferase inhibitory activity, Journal of Biological Chemistry 265(14) (1990) 8042–8051.

[100] G.F. Barnard, S.K. Erickson, A.D. Cooper, Regulation of lipoprotein receptors on rat hepatomas in vivo, Biochimica et Biophysica Acta (BBA)-Lipids and Lipid Metabolism 879(3) (1986) 301–312.

[101] R.C. Brown, M.L. Blank, J.A. Kostyu, P. Osburn, A. Kilgore, F. Snyder, Analysis of tumor-associated alkyldiacylglycerols and other lipids during radiation-induced thymic leukemogenesis, Proceedings of the Society for Experimental Biology and Medicine 149(3) (1975) 808–813.

[102] S.K. Erickson, A.D. Cooper, G.F. Barnard, C.M. Havel, J.A. Watson, K.R. Feingold, A.H. Moser, M. Hughes-Fulford, M.D. Siperstein, Regulation of cholesterol metabolism in a slow-growing hepatoma in vivo, Biochimica et Biophysica Acta (BBA)-Lipids and Lipid Metabolism 960(2) (1988) 131–138.

[103] H. Konishi, H. Okajima, Y. Okada, H. Yamamoto, K. Fukai, H. Watanabe, High levels of cholesteryl esters, progesterone and estradiol in the testis of aging male Fischer 344 rats: feminizing Leydig cell tumors, Chemical and pharmaceutical bulletin 39(2) (1991) 501–504.

[104] J.M. Olsson, L.C. Eriksson, G. Dallner, Lipid compositions of intracellular membranes isolated from rat liver nodules in Wistar rats, Cancer research 51(14) (1991) 3774–3780.

[105] S. Ruggieri, A. Fallani, Lipid composition of Yoshida ascites hepatoma and of livers and blood plasma from host and normal rats, Lipids 14(4) (1979) 323–333.

[106] S. Ruggieri, A. Fallani, D. Tombaccini, Effect of essential fatty acid deficiency on the lipid composition of the Yoshida ascites hepatoma (AH 130) and of the liver and blood plasma from host and normal rats, Journal of lipid research 17(5) (1976) 456–466.

[107] D.J. Talley, J.A. Sadowski, S.A. Boler, J.J. Li, Changes in lipid profiles of estrogen-induced and transplanted renal carcinomas in Syrian hamsters, International journal of cancer 32(5) (1983) 617–621.

[108] R. Wood, F. Chumbler, R.D. Wiegand, Effect of dietary cyclopropene fatty acids on the octadecenoates of individual lipid classes of rat liver and hepatoma, Lipids 13(4) (1978) 232–238.

[109] H. Xu, H. Xia, S. Zhou, Q. Tang, F. Bi, Cholesterol activates the Wnt/PCP-YAP signaling in SOAT1-targeted treatment of colon cancer, Cell death discovery 7(1) (2021) 1–13.

